# Predicting Suicide Risk in Bipolar Disorder patients from Lymphoblastoid Cell Lines genetic signatures

**DOI:** 10.1101/2024.05.30.596645

**Authors:** Omveer Sharma, Ritu Nayak, Liron Mizrahi, Wote Amelo Rike, Ashwani Choudhary, Yara Hussein, Idan Rosh, Utkarsh Tripathi, Aviram Shemen, Alessio Squassina, Martin Alda, Shani Stern

## Abstract

This research investigates the genetic signatures associated with a high risk of suicide in Bipolar disorder (BD) patients through RNA sequencing analysis of lymphoblastoid cell lines (LCLs). By identifying differentially expressed genes (DEGs) and their enrichment in pathways and disease associations, we uncover insights into the molecular mechanisms underlying suicidal behavior. LCL gene expression analysis reveals significant enrichment in pathways related to primary immunodeficiency, ion channel, and cardiovascular defects. Notably, genes such as *LCK*, *KCNN2*, and *GRIA1* emerged as pivotal in these pathways, suggesting their potential roles as biomarkers. Machine learning models trained on a subset of the patients and then tested on other patients demonstrate high accuracy in distinguishing low and high-risk of suicide in BD patients. Moreover, the study explores the genetic overlap between suicide-related genes and several psychiatric disorders. This comprehensive approach enhances our understanding of the complex interplay between genetics and suicidal behavior, laying the groundwork for future prevention strategies.

## 1 Introduction

Suicide is a deeply complex and tragic phenomenon that continues to be a significant public health concern worldwide. Annually, the global incidence of suicide surpasses 700,000 fatalities exhibiting variations observed among distinct age cohorts and geographical regions [1, 2]. Suicide often unfolds within the framework of psychiatric work [3]. Research findings show that more than 90% of individuals who succumb to suicide have a diagnosed psychiatric illness and most individuals who attempt suicide also experience such disorders [4, 5]. A significant number of individuals diagnosed with BD may undergo at least one episode of self-harm in their lifetime, while approximately 20% died by suicide [6, 7]. Thus, BD carries the highest risk of suicidal behavior among other psychiatric disorders [8–10] and currently, there is no way to identify those BD patients that are at a high risk of suicide.

The risk associated with suicide in BD varies depending on the disease attributes and the phase or stage of the illness. Existing studies suggest that BD II has a higher suicide rate than BD I. This observation emphasizes the multifaceted severity of BD II [11] with major depressive episodes, carrying the highest risk of suicide among individuals with BD [12]. Following closely are BD patients with mixed episodes, where both depression and mania are present, posing a significant risk as well [13]. Conversely, during manic episodes, marked by elevated mood and energy levels, the patient has the lowest risk of suicide [14]. Additionally, individuals with rapid cycling BD face an increased risk compared to those without rapid cycling patterns [15]. When considering sex differences, certain studies suggest that men exhibit higher rates of fatal suicide attempts, whereas women have more non-fatal suicide attempts[16, 17]. However other research indicates no sex disparity in suicide attempts [18].

In patients with BD, a history of prior suicide attempts emerges as one of the most robust predictors of both fatal and non-fatal suicide attempts [19, 20]. While a significant portion, approximately 56% of those who die by suicide do so on their first attempt [19, 21] the majority of individuals who attempt suicide non-fatally do not ultimately die by suicide [22]. Subsequently, a few studies have also highlighted the strong association between a first-degree family history of suicide and suicide deaths among individuals with BD [23, 24]. Thus, it confirms the presence of both a family history and a personal history of suicide attempts as a critical suicide risk factor in BD [14, 25]. Other factors include early life trauma such as childhood abuse or stress, which are linked to higher rates of suicide attempts in individuals with BD [26–28]. Moreover, the combination of childhood abuse and drug usage drastically increases the likelihood of suicide attempts [29]. Psychosocial stressors, including interpersonal conflicts and occupational issues, have also been associated with increased suicide risks [30, 31].

The study of BD has been revolutionized by various cellular model systems, especially using the induced pluripotent stem cell (iPSC) model which allows researchers to explore the cellular and molecular mechanisms underlying the disorder in neuronal cell types derived from patients themselves. The iPSC-based neuronal models from BD patients have reported mitochondrial dysfunction, neuron hyper-excitability, disruption of calcium signaling etc [32, 33]. These BD studies revealed key electrophysiological differences in hippocampal neurons of BD patients, particularly between those responsive (LR) and non-responsive (NR) to lithium treatment. An elevated fast after-hyper-polarization (AHP) was shown to be a distinct trait in BD neurons, which has been implicated in various neurological disorders [34–36]. The study led to the development of computational models for the prediction of lithium treatment [37, 38]. Given the high risk of suicide among patients with BD, there is a strong need to understand the neurobiological and genetic factors of suicide, an iPSC model may shed more light on the related mechanism [39].

Genome-wide association studies (GWAS) have paved the way in identifying genetic loci associated with suicide across psychiatric disorders [40, 41]. Notable genes including *5-HTT*, *SLC6A4*, and the *5-HT1* to *5-HT7* series involved in serotonergic neurotransmission, along with tryptophan hydroxylase genes *TPH1* and *TPH2*, which are identified for their crucial roles in the neurotransmission dysfunctions tied to various suicidal behaviors [42–46]. Research also extends to genes such as *BDNF* and its receptor *NTRK2*, and others like *COMT* and *MAPK1*, exploring their links to suicide risk [47–52]. Furthermore, genetic variations in genes like *AKT1P*, *GSK3B*, and the Alpha2a-adrenergic receptor (*ADRA2*) are associated with impulsivity, a significant risk factor in BD suicide [53–56]. Additionally, the *PENK* gene, along with *IL-7* and *TMX3*, involved in stress and anxiety responses, were linked to suicidal behavior in BD patients [57]. Comprehensive multi-ancestry GWAS analysis has pinpointed genes such as *DRD2*, *FURIN*, *NLGN1*, *SOX5*, *PDE4B*, and *CACNG2* as involved in suicidal behavior, alongside identifying potential biomarkers within the immunoglobulin gene family for BD [38, 58]. Despite these advancements, a classification based solely on GWAS data remains challenging.

In this study, we have sought biomarkers for suicidal tendencies in BD patients to construct machine-learning algorithms that will enable a good prediction of the patients who are at risk of attempting suicide. To do this, we have grown LCLs from 20 patients and extracted their RNA. Six of the patients died by suicide; blood samples were collected before their deaths. Seven BD patients have been monitored over a long period and have not attempted suicide over many years. The remaining 7 patients have either attempted suicide or have a family history of suicide (and one did not have either). Interestingly, when analyzing enrichment for biological pathways, we found brain-related and psychiatric disease-related pathways, despite the source of the RNA being from LCLs. The patients were then divided into two groups of those who died by suicide and those who were at a low risk since they were monitored over the years. We applied machine learning (ML) algorithms to train on DEGs derived from the RNA sequencing data. Using cross-validation schemes, we were able to show that prediction is possible with a very low error rate using RNA sequencing data of the patient’s LCLs.

The structure of this work is as follows: Section II presents the methodology. Experimental results are discussed in Section III. Section IV elaborates on the discussion, while Section V provides the conclusion.

## 2 Methodology

### 2.1 Ethics Statement and LCLs Collection

All participants provided informed consent before participating in the study. Approval for the study was obtained from the Research Ethics Board of the University of Cagliari, Italy. The participants, diagnosed with BD, were categorized into subtypes based on those patients who have died by suicide and those who are at a low risk of suicide.

#### 2.1.1 Data set

We conducted RNA sequencing on samples from 20 patients diagnosed with BD. Of these, the status of 13 patients—whether they have died by suicide or have tried to attempt suicide was known. Among these 13, six patients died by suicide, while the remaining seven were monitored over the years and had no family history of suicide nor have they attempted suicide. Further details on these patients can be found in Table 1. For this study, the RNA sequencing data from these 13 patients were used to train and test machine learning models. Additional information about the samples will be available in the GitHub file at this link (https://github.com/omveersharmanet/Predicting-Suicide-Risk-in-Bipolar-Disorder-patients.git).

**Table 1:**
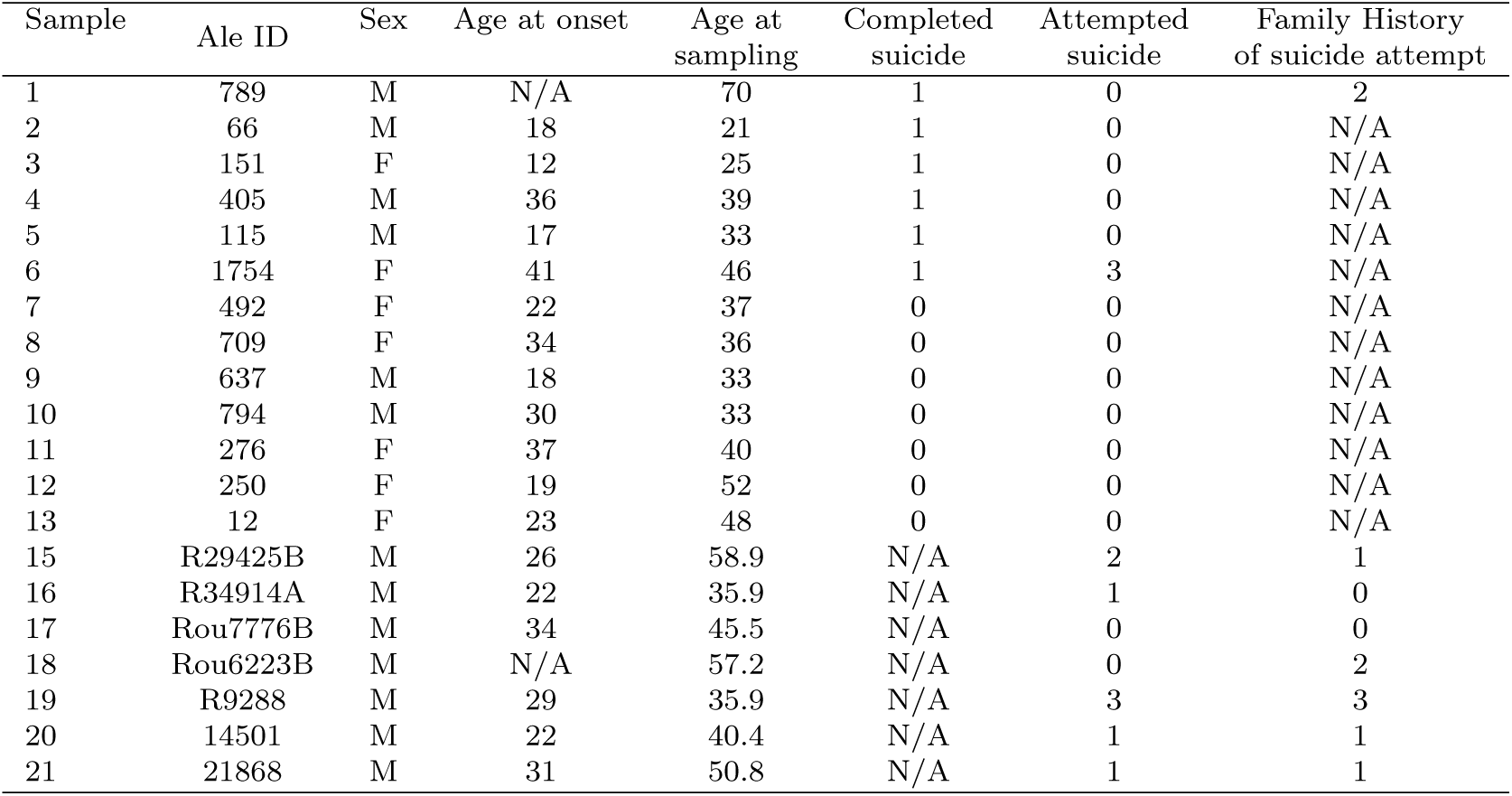
Details of BD patients add family history.

| Sample | Ale ID | Sex | Age at onset | Age at sampling | Completed suicide | Attempted suicide | Family History of suicide attempt |
| --- | --- | --- | --- | --- | --- | --- | --- |
| 1 | 789 | M | N/A | 70 | 1 | 0 | 2 |
| 2 | 66 | M | 18 | 21 | 1 | 0 | N/A |
| 3 | 151 | F | 12 | 25 | 1 | 0 | N/A |
| 4 | 405 | M | 36 | 39 | 1 | 0 | N/A |
| 5 | 115 | M | 17 | 33 | 1 | 0 | N/A |
| 6 | 1754 | F | 41 | 46 | 1 | 3 | N/A |
| 7 | 492 | F | 22 | 37 | 0 | 0 | N/A |
| 8 | 709 | F | 34 | 36 | 0 | 0 | N/A |
| 9 | 637 | M | 18 | 33 | 0 | 0 | N/A |
| 10 | 794 | M | 30 | 33 | 0 | 0 | N/A |
| 11 | 276 | F | 37 | 40 | 0 | 0 | N/A |
| 12 | 250 | F | 19 | 52 | 0 | 0 | N/A |
| 13 | 12 | F | 23 | 48 | 0 | 0 | N/A |
| 15 | R29425B | M | 26 | 58.9 | N/A | 2 | 1 |
| 16 | R34914A | M | 22 | 35.9 | N/A | 1 | 0 |
| 17 | Rou7776B | M | 34 | 45.5 | N/A | 0 | 0 |
| 18 | Rou6223B | M | N/A | 57.2 | N/A | 0 | 2 |
| 19 | R9288 | M | 29 | 35.9 | N/A | 3 | 3 |
| 20 | 14501 | M | 22 | 40.4 | N/A | 1 | 1 |
| 21 | 21868 | M | 31 | 50.8 | N/A | 1 | 1 |

### 2.2 LCL Growth and Expansion

We received LCLs derived from Peripheral blood mononuclear cells (PBMCs) from both groups of patients. Briefly, PBMCs were isolated from blood samples using BD vacutainer according to the manufacturer’s guidelines. Following which they were exposed to Epstein-Barr virus (EBV) harvested from the B95-8 cell line. Within a week post-infection, the cells transformed and began to aggregate, forming large clusters over time.

The LCLs were cultured in T25 tissue culture flasks, using a complete RPMI medium composed of RPMI 1640 (Biological Industries, Cat no: 01-100-1A), 1X Anti-Anti (Thermofisher Scientific, Cat no: 15240062), 1% Glutamax (Thermofisher Scientific, Cat no: 35050061), 1% Sodium pyruvate (Thermofisher Scientific, 11360070), and 15% heat-inactivated fetal bovine serum (Sigma, Cat no: F9665), as previously outlined [59]. Media changes were performed every other day. Passaging of the LCL was performed to maintain optimal cell growth and prevent over confluency. Cells were sub cultured when reaching a density of 200,000 cells/ml to sustain a logarithmic growth.

#### 2.2.1 RNA Extraction, Sequencing, and Analyses

Total cellular RNA was extracted from 5 million LCLs using 800µl of TRIzolTM reagent (Thermofisher Scientific, Cat no. 15596026) and subsequently chilled on ice for 5 minutes before long-term storage at -80 *^◦^* Celsius. The total RNA was carefully isolated using the zymo RNA clean & concentrator kit, according to manufacturer’s protocol. Following the extraction, the quality and integrity of the RNA samples were quantified using ND-1000 Nanodrop spectrophotometer (Thermo Sci). All RNA samples displayed an RNA integrity number (RIN) within 7-8, confirming both good quality and suitability for sequencing.

RNA sequencing and analysis Libraries of RNAs extracted from LCLs derived from BD patients were prepared using the TruSeq RNA Library Prep Kit v2 (Illumina), adhering to the manufacturer’s guidelines. Quality control was conducted on the raw FASTQ files using FastQC (v0.11.5) [60]. Reads were aligned to the human genome (GRCh38.104) and quantified with STAR (v2.7.9a) [61]. Differential gene expression analysis was carried out using DESeq2 (v1.34.0) [62]. To mitigate false positives in identifying DEGs, a false discovery rate (FDR) analysis was applied using the Benjamini-Hochberg (BH) procedure to adjust p-values for multiple hypothesis testing [63]. Genes were considered significant DEGs if they met the criteria of a p-value *<* 0.05 and a *log*_2_ fold-change *≥ |*0.58*|*, with a total of 843 genes meeting these thresholds. The step-by-step process for identifying DEGs is illustrated in Fig. 1(a). The top 50 genes were selected as predictors for further machine-learning analysis.

**Fig. 1:**
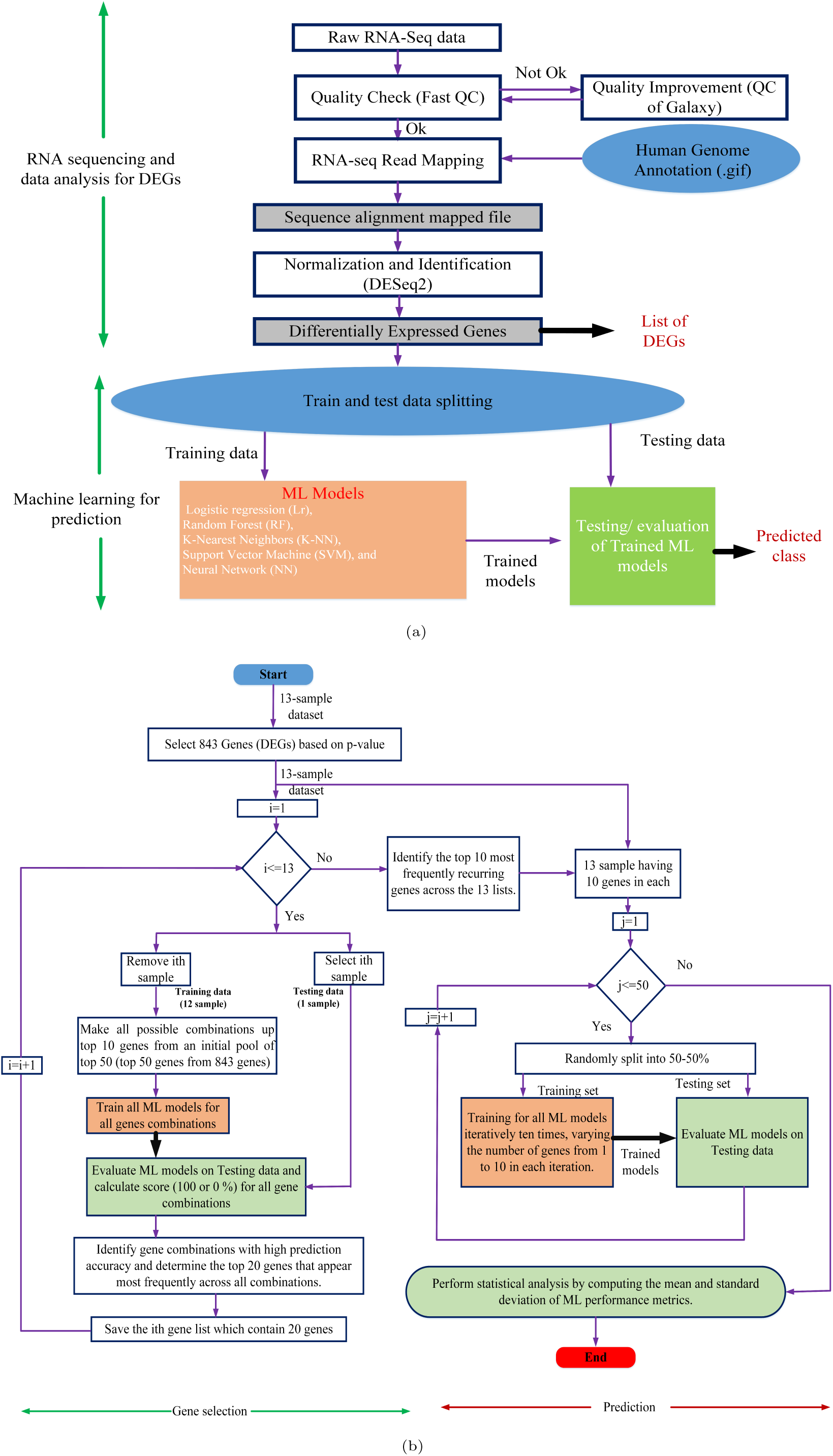
(a) A Flowchart illustrating the process of identifying differentially expressed genes (DEGs) and training and testing the machine learning models. (b) A step-by-step training and testing of the machine learning models.

### 2.3 Machine Learning Predictor Analysis

We assessed five distinct supervised classifiers for their ability to predict suicidal and non-suicidal outcomes; Logistic Regression (LR) is a fundamental statistical model utilized for binary classification, leveraging a logistic function to estimate probabilities and optimize parameters through maximizing data likelihood, though its performance may falter with non-linear data relationships. The K-Nearest Neighbors (K-NN) algorithm, a non-parametric method, classifies new instances by using the majority vote from the ’K’ nearest training samples, but struggles with large or high-dimensional datasets due to the heavy computational demand of distance calculations. Support Vector Machine (SVM) creates a hyperplane in high-dimensional space to separate classes with optimal margins. It adapts well to nonlinear boundaries using the kernel trick, although tuning its hyperparameters can be challenging and resource-intensive [64, 65]. Random Forest (RF) enhances robustness and accuracy by building multiple decision trees and integrating their outcomes [66], which are suitable for various tasks but potentially complex and difficult to interpret. In our previous research, we discovered that SVM and RF were the most effective machine-learning algorithms in distinguishing between lithium responders and non-responders among BD patients [37, 67–69]. Inspired by the human brain, neural Networks (NN) excel in pattern recognition and learning complex nonlinear relationships, making them powerful yet resource-heavy and challenging to manage due to their propensity for over-fitting [70]. Each algorithm offers unique strengths and limitations, with the choice often depending on data characteristics and specific problem needs. Each of these classifiers has been carefully selected to address different aspects of the predictor analysis in our study, taking into account their respective strengths and limitations according to the specific requirements of the data and the task at hand. Logistic regression, random forest, and neural network classifiers gave the best performance with high accuracy (*>*96%).

The feature selection process (predictors/genes) is illustrated in Fig. 1(b). We have employed an intensive approach that combines leave-one-out cross-validation (LOOCV) with an exhaustive feature selection process to identify the most crucial features for our model, especially useful for smaller dataset. Specifically, we utilized LOOCV for our dataset, which consists of 13 samples. This method involved training our model on 12 samples and testing on the remaining one, cycling through iteratively so that each sample served as the test set once, resulting in 13 total iterations. Concurrently, during each LOOCV iteration, we examined all possible combinations of up to 10 genes from the initial top 50 DEGs, assessing the LR model’s accuracy with these subsets on the left-out sample, where the outcome was binary—100% or 0% accuracy based on correct or incorrect predictions. Successful combinations led to a tally of each gene’s appearance frequency, and after 13 iterations, the top 20 genes were identified based on this frequency. This process was repeated to yield 13 lists of the top 20 genes, from which we further distilled the data to pinpoint the top 10 consistently appearing genes. Finally, to evaluate our ML models, these top 10 genes were used to split the full dataset into training and testing sets multiple times (50 splits with a 50-50% ratio), and the mean accuracy from these tests provided a robust assessment of the model’s effectiveness.

The effectiveness of the approach was validated by testing five supervised classification algorithms using the specified input features for binary classification. The outcomes of the model predictions were classified into four categories: true positives (tp), false positives (fp), true negatives (tn), and false negatives (fn). Evaluation metrics included accuracy, the Receiver Operating Characteristic (ROC) curve, and the confusion matrix. To ensure robustness, cross-validation methods were applied, with results being averaged or aggregated for the confusion matrix across iterations. Performance was assessed using the Area Under the Curve (AUC) of the ROC and metrics from the confusion matrix such as Accuracy and Precision. Fig. 1(b) illustrates the feature selection process (predictors/genes) and the training and testing of machine learning models using the selected genes. Accuracy of ROC:

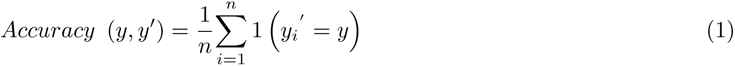

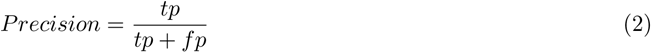

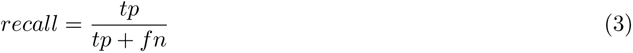

Where, *y^′^* represents the model’s prediction, while *y* denotes the actual class.

## 3 Results

Our research focused on developing biomarkers that are easily and cheaply extracted from biological samples that will aid in identifying BD patients who are at an increased risk of suicide. For this, we have grown LCLs of BD patients who died by suicide (6 samples) and BD patients who have been followed over many years and have not attempted suicide, nor have any family history of suicide. For this, we performed the following steps:

1) Identifying DEGs between patients who died by suicide (‘SUICIDE’) compared to those that are at a low risk (no attempts over many years and no family history ‘NON-SUICIDE’),
2) selecting the most predictive genes, and
3) training and testing machine learning models first on these two groups and then on another set of patients who are at an increased risk of suicide, as shown in Fig. 1(b).

### 3.1 The Identification of DEGs in LCLs of BD Patients (‘SUICIDE’ and ‘NON-SUICIDE’ Groups)

To identify biomarkers for suicide risk, we performed sequencing of RNA extracted from LCLs of 20 BD patients. Of these 6 patients died by suicide (‘SUICIDE’), 7 have been followed over the years and have never attempted suicide, nor do they have a family history of suicide (‘NON-SUICIDE’), and the classification of the remaining 7 patients for whom we have a family history of suicide and some have attempted non-fatal suicide (‘UNKNOWN’). Differential gene expression analysis was conducted between the first two groups of BD subjects (‘SUICIDE’ and ‘NON-SUICIDE’), as detailed in the Methods section (2). We identified 843 DEGs with a fold change greater than 1.5 (*log*_2_ fold-change *≥ |*0.58*|*) and a p-value of less than 0.05. These are presented in Fig. 2(b) as a heatmap. Fig. 2(b) displays the gene counts for the 30 most significantly DEGs based on the lowest p-values. DEGs between ’SUICIDE’ and ’NON-SUICIDE’ LCLs exhibited enrichment in pathways related to “Primary immunodeficiency” such as *LCK*, *CD8A*, *AICDA*, *ZAP70*, and *CIITA*. Specifically, *LCK*, *AICDA*, *ZAP70*, and *CIITA* demonstrate higher gene counts in ’SUICIDE’ LCLs compared to ’NON-SUICIDE’ LCLs, indicating increased expression levels associated with suicidal behaviour. However, *CD8A* shows a contrasting pattern, with higher gene counts observed in ’NON-SUICIDE’ LCLs compared to ’SUICIDE’ LCLs, suggesting a distinct expression profile in non-suicidal individuals. DEGs between the ‘SUICIDE’ and ‘NON-SUICIDE’ groups were enriched in Immunoglobulin genes, such as *IGHV1-3*, *IGHV4-31*, *IGHV3-33*, *IGHV1-46*, *IGHV4-59*, *IGHV4-61* were enriched in ’NON-SUICIDE’ samples whereas *IGKV4-1* was enriched in ’SUICIDE’ samples. 0.8% (470 genes) of the total human genes are immunoglobulin genes. However, among the DEGs, 2.3% (19 out of 843 DEGs) are immunoglobulin genes. Furthermore, when examining DEGs with minimum counts of 10, 100, and 1000, the proportion of immunoglobulin-related genes increases to 2.3%, 3.1%, and 5.2%, respectively. This pattern indicates a significant enrichment of immunoglobulin-related genes in the DEGs, as illustrated in Supplementary Fig. 1(a).

**Fig. 2:**
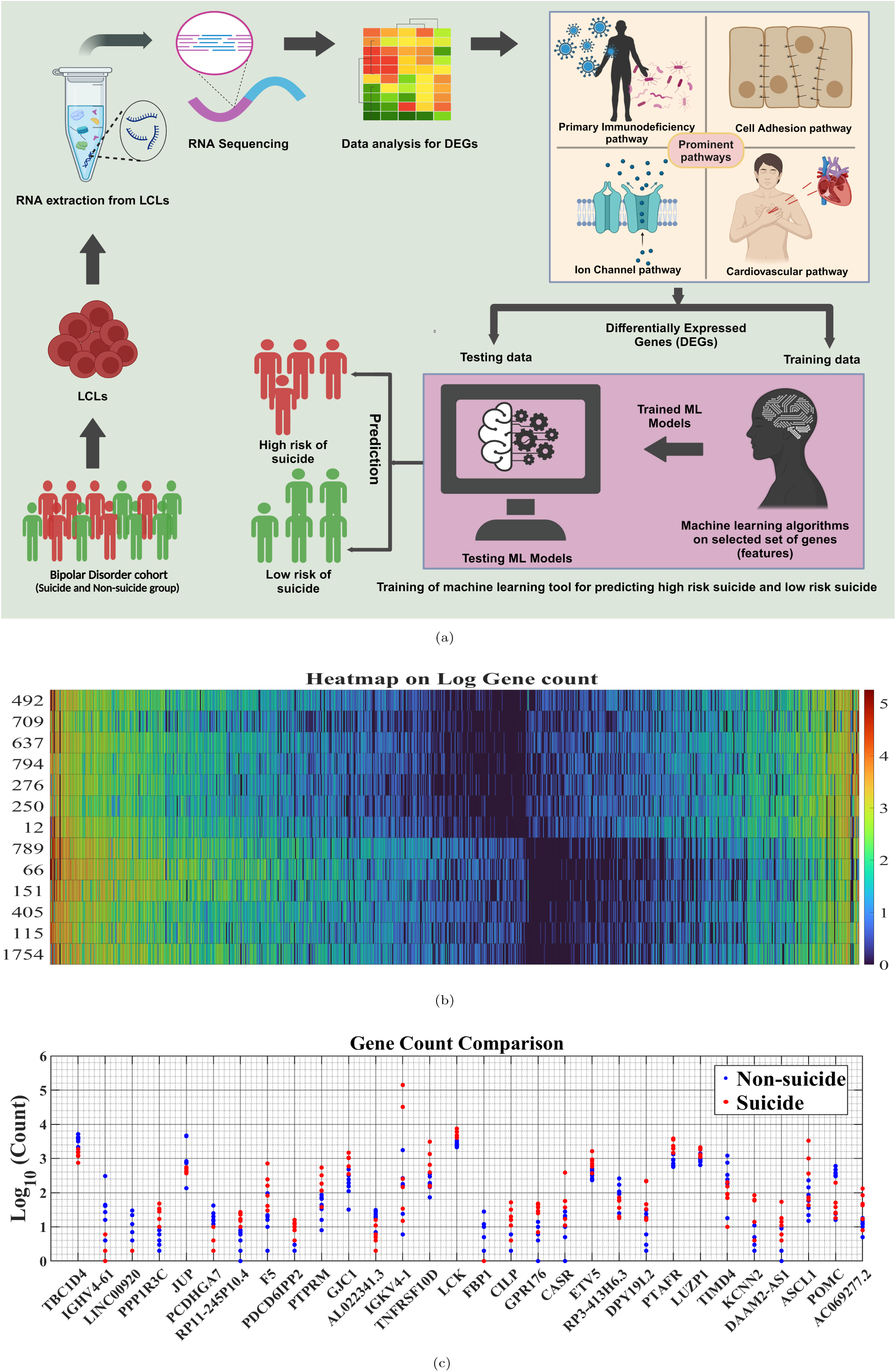
Flow chart, Genes and pathways that distinguish the two groups of ‘SUICIDE’ and ‘NON-SUICIDE’. (a) Flow chart (b) A heatmap of log10 gene count of the DEGS between the two groups of ‘SUICIDE’ and ‘NON-SUICIDE’. The heat map was organized based on the descending order of the subtraction of the average read count of the two groups (‘SUICIDE’-‘NON-SUICIDE’). (c) The gene count of the top 30 genes with the lowest p-values is displayed, with red indicating suicidal individuals and blue indicating non-suicidal individuals.

### 3.2 Associated dysregulated pathways and diseases in LCLs of BD patients (‘SUICIDE’ vs. ‘NON-SUICIDE’)

Further analysis of the DEGs included exploring KEGG pathways, disease associations, and functional annotations, as depicted in Fig. 3. The KEGG pathway analysis highlights the intricate biochemical and physiological networks potentially involved in the underlying conditions studied, as shown in Fig. 3(a). For example, pathways such as primary immunodeficiency, phenylalanine, tyrosine, tryptophan biosynthesis, and hematopoietic lineage are significantly dysregulated between BD patients from the two groups (‘SUICIDE’ vs. ‘NON-SUICIDE’). Additionally, the Intestinal Immune Network for IgA Production and Primary Immunodeficiency pathways suggest a significant role of immune system functions in the pathology, potentially linking systemic immune responses to neurological and psychiatric conditions.

**Fig. 3:**
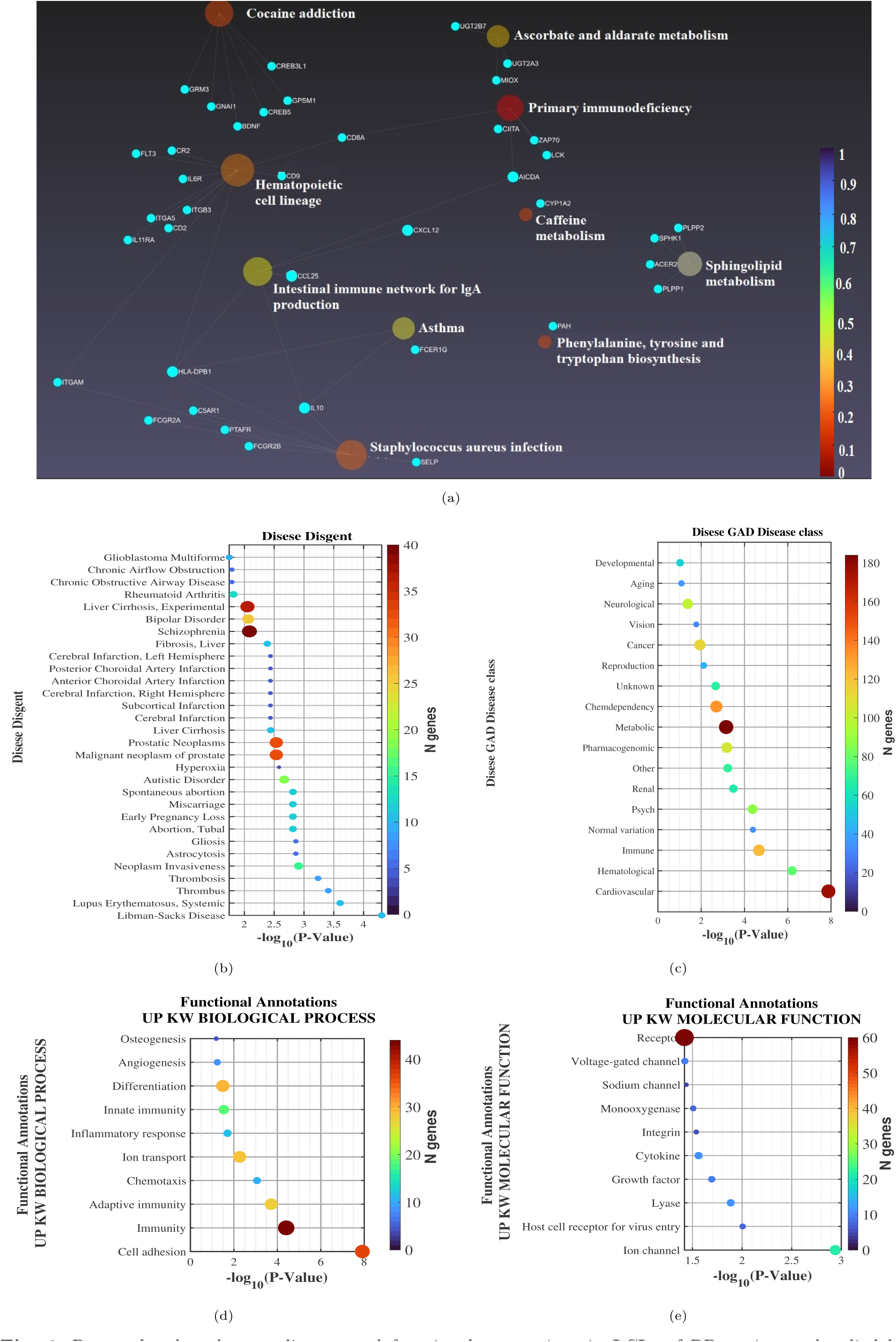
Dysregulated pathways, disease, and functional annotations in LCLs of BD patients who died by suicide compared patients at a low risk of suicide. (a) A diagram displaying the dysregulated KEGG pathways associated with the DEGs between the two groups. (b) The most prevalent conditions in the DisGeNET database, such as Libman-Sacks Disease, Systemic Lupus Erythematosus, Thrombosis, Neoplasm Invasiveness, Gliosis, Astrocytosis, and various conditions related to pregnancy loss and disorders. (c) Prominent GAD disease classes, including Cardiovascular, Hematological, Immune, Psychiatric, Renal, and others. (d) Significant biological processes like Cell Adhesion, Immunity, Chemotaxis, and Inflammatory Response. (e) Molecular functions associated with the DEGs, including Ion Channels and Growth Factors.

In the Disease DisGeNET analysis, a wide array of conditions linked to the DEGs was identified, indicating a broad impact of these genetic variations across various diseases, as shown Fig. 3(b). The list includes autoimmune and inflammatory diseases such as Libman-Sacks Disease and Systemic Lupus Erythematosus, as well as conditions related to abnormal blood clotting, like thrombosis. Importantly and surprisingly, many brain-related pathways came up in the analysis, despite the origin of the cells (LCLs). These pathways include BD Schizophrenia, Cerebral infraction Left and Right hemispheres, Subcortical infraction, Autistic Disorder, neuroinflammatory diseases like Gliosis, Astrocytosis, and more.

The Disease GAD (Genetic Association Database) list encompasses a wide range of disorders, highlighting the complex interplay between genetics and various health conditions, as shown in supplementary Fig. 1(b). Diseases such as Hypertension, Asthma, and Coronary Artery Disease reflect chronic conditions that significantly impact public health and are influenced by both genetic predispositions and lifestyle factors. Autoimmune and inflammatory diseases like Systemic Lupus Erythematosus and Rheumatoid Arthritis, as well as metabolic disorders such as Type 2 Diabetes, underscore the genetic basis of immune system dysfunction and metabolic dysregulation. Furthermore, the inclusion of conditions like schizophrenia and substance use disorder illustrates the genetic factors contributing to psychiatric disorders and addictive behaviors, demonstrating the broad spectrum of diseases that genetic research can potentially address. In our research, we conducted an in-depth analysis of diseases characterized by specific up-regulated genetic or molecular markers, focusing on how these enhancements influence disease pathology, as shown in supplicatory Fig. 1(b). This exploration has been pivotal in identifying key biological pathways and potential therapeutic targets, enhancing our understanding of their underlying mechanisms.

The GAD Disease class encompasses a broad spectrum of medical categories, reflecting the diverse genetic underpinnings and environmental influences on various health conditions, as shown in Fig. 3(c). Interestingly, our data indicates that neurological and developmental pathways are dysregulated even in the white blood cells of the patients. Another interesting pathway that was dysregulated was “AGING” as accelerated aging was reported in BD [71] and according to our data, this may be even further implied in BD patients who died by suicide. “PSYCH” was one of the top dysregulated pathways in the LCLs. The “IMMUNE” pathway is among the top 3 dysregulated pathways. Immune pathways have been shown repeatedly to be dysregulated in BD [72]. The most dysregulated pathway was “CARDIOVASCULAR”. Associations between BD and heart defects were previously shown [73]. Our data shows that these pathways are even more dysregulated in BD patients, who are at a high risk of suicide.

We also conducted a detailed analysis across five functional annotation categories for whom the DEGs between the groups were enriched including UP KW BIOLOGICAL PROCESS, UP KW CELLULAR COMPONENT, UP KW MOLECULAR FUNCTION, UP KW PTM, and UP SEQ FEATURE as shown in Fig. 3(d)&3(e) and Supplementary Fig. 1 (c, d, and e). Specifically, within the UP KW BIOLOGICAL PROCESS category, we identified significant dysregulation in processes such as cell adhesion, which is vital for cellular assembly and integrity, and various immune-related functions, including adaptive and innate immunity, underscoring the genes’ pivotal roles in systemic defence mechanisms, as shown in Fig. 3(d). Additionally, this category highlighted the involvement of genes in essential developmental and reparative processes such as angiogenesis and osteogenesis, crucial for maintaining vascular and skeletal health. In our study, the UP KW MOLECULAR FUNCTION revealed critical roles of various proteins in cellular mechanisms, with a particular emphasis on ion channels, sodium channels, and voltage-gated channels, which are essential for electrical signalling and cellular communication and have been shown to alter in neurons derived from BD patients[74], as shown in Fig. 3(e). Proteins such as cytokines and growth factors were also identified as changed in LCLs of suicide victims. These proteins are important for cell signalling and regulatory processes that influence cell growth, differentiation, and immune responses, suggesting implications for suicide. Additionally, the identification of host cell receptors for virus entry, integrins, and receptors underscores the relevance of cell-environment interactions in suicide, which facilitates crucial processes such as viral entry, cellular adhesion, and signal transduction across cellular membranes.

### 3.3 Selecting the Most Predictor Genes

We initiated this study with the goal of developing biomarkers that could be quickly and easily implemented in clinical settings. Consequently, it was essential to establish a robust protocol that facilitates the prediction of BD patients who are at a high risk of suicide using RNA-seq datasets, as illustrated in Fig. 2(a). As described in the methodology section (2), the top 50 genes were selected from a pool of 843 DEGs to serve as predictors for machine learning algorithms. We implemented a rigorous approach to identify the most dominant and predictive genes using LOOCV and exhaustive feature selection, well-suited for our dataset. Our method involved training the model on 12 samples and testing on the remaining one, cycling through each sample as the test set, completing 13 iterations. During each iteration, we tested all combinations of up to 10 genes from the initial 50, assessing the Logistic Regression model’s accuracy on the left-out sample, which was binary—either 100% or 0% based on prediction accuracy. This allowed us to identify the top 20 genes appearing most frequently across iterations. After repeating this process, we narrowed down to the top 50 genes consistently identified across the 13 lists. The gene counts and heat maps for these top 50 genes are shown in Figs. 4(a) & 4(b). We further narrowed down the top 10 genes to train and test all ML models, as shown in Fig. 1(b).

**Fig. 4:**
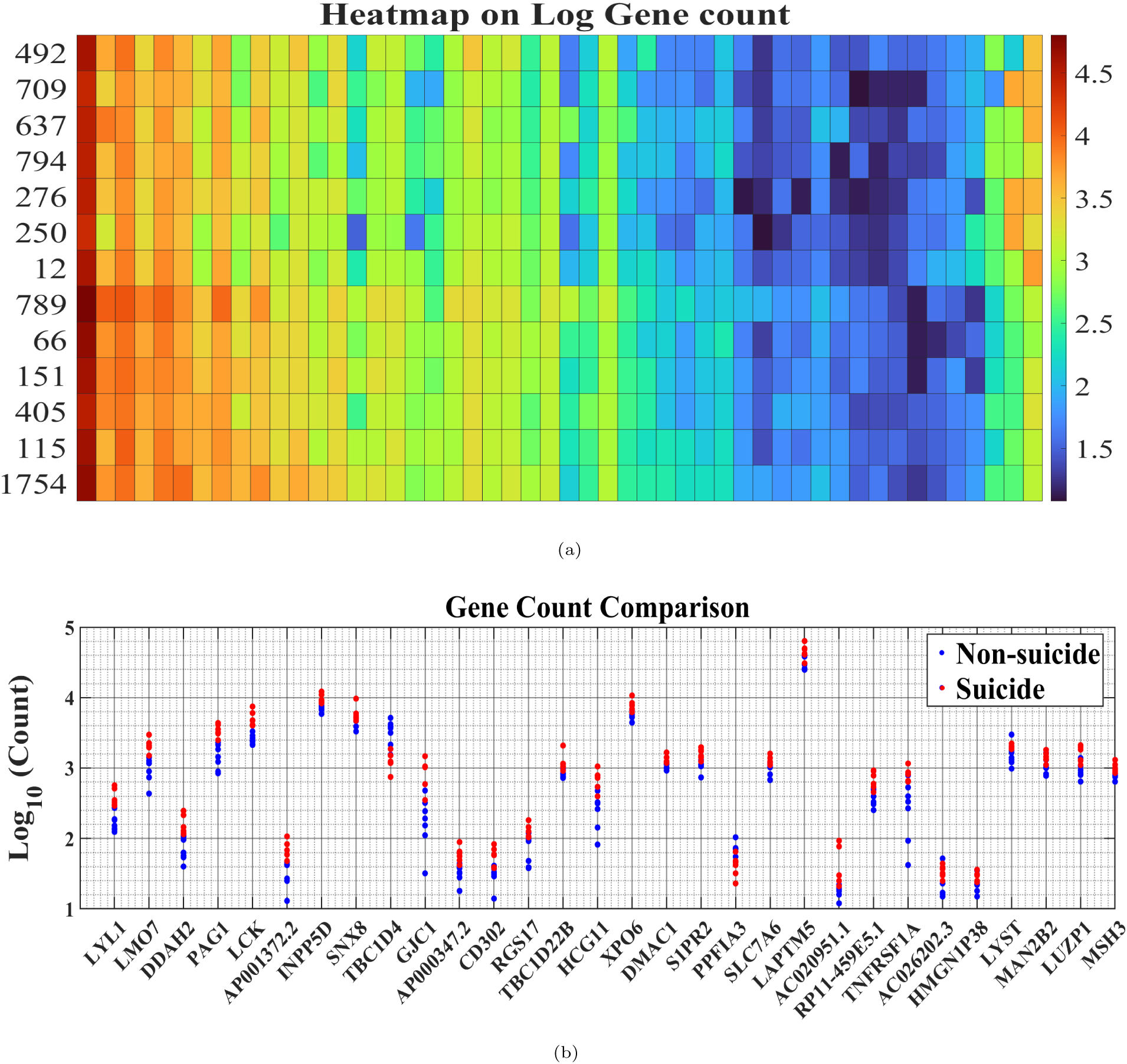
Top predictive genes for suicide risk. (a) A heat map, the heat map was organized based on the descending order of the subtraction of the average read count of the two groups (‘SUICIDE’-‘NON-SUICIDE’). (b) Gene counts for the most dominant predictor for ML.

### 3.4 Training and Testing of Machine-learning Models

After identifying the top 10 genes (*LYL1*, *LMO7*, *DDAH2*, *PAG1*, *LCK*, *AP001372.2*, *INPP5D*, *SNX8*, *TBC1D4*, *GJC1*), we proceeded to train and test various machine learning (ML) models. We employed five types of supervised classification algorithms: Logistic Regression (LR), Random Forest (RF), K-Nearest Neighbors (K-NN), Support Vector Machine (SVM), and Neural Network (NN). To avoid over-fitting, we used cross-validation methods, as detailed in the Methods section (2) (refer to Fig. 1(b)). We trained and tested the models using subsets varying from 1 to 10 predictors, as illustrated in Fig. 5(a). Notably, we found that 8 features provided high accuracy of more than 95%.

**Fig. 5:**
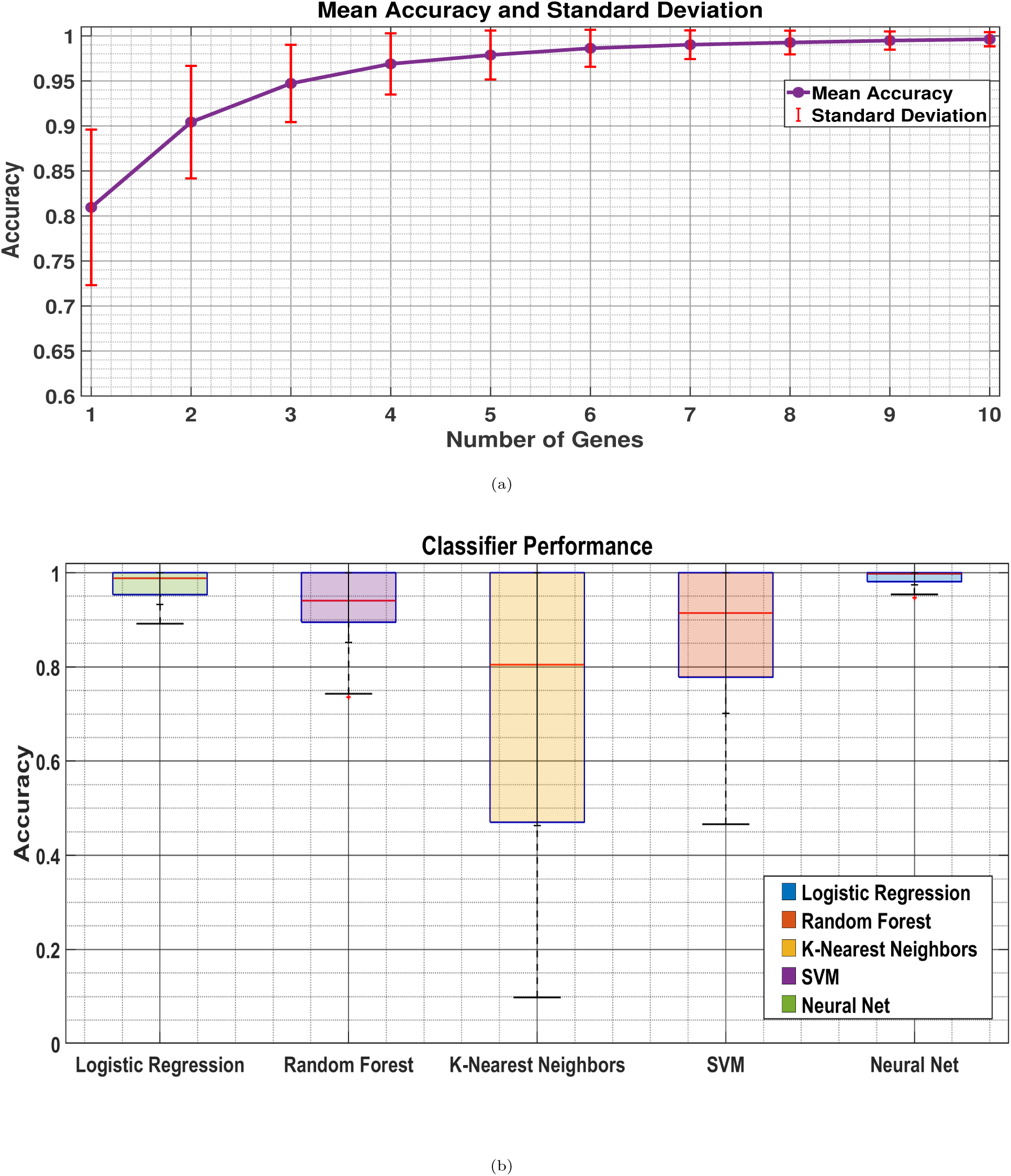
Prediction accuracy by varying the number of genes. (a) The variation in model accuracy for a logistic regression machine learning (ML) model in relation to the number of genes used as predictor features, ranging from 1 to 10 genes. (b) The accuracy of various ML models when specifically using 8 predictors.

We narrowed down to 8 genes with high predictive power of suicidal and non-suicidal outcomes, as indicated in Fig. 5b. Here, we divided the data with 50% for training and 50% for testing, a process repeated 50 times to ensure the robustness of our ML algorithms. The accuracy results were as follows: Logistic Regression: 98.3%*±*5.0%, Random Forest: 96.0%*±*10.8%, K-Nearest Neighbors: 77.0%*±*30.7%, SVM: 90.7%*±*20.6%, and Neural Network: 99.7%*±*2.3%. Both Logistic Regression and Neural Network showed higher accuracy rates compared to the other ML models. Fig. 6 displays the performance of these classifiers through the area under the ROC curves and confusion matrices, with the following accuracies: Logistic Regression: 0.89*±*0.00, Random Forest: 0.99*±*0.00, K-Nearest Neighbors: 0.99*±*0.00, SVM: 0.61*±*0.22, and Neural Network: 0.67*±*0.20.

**Fig. 6:**
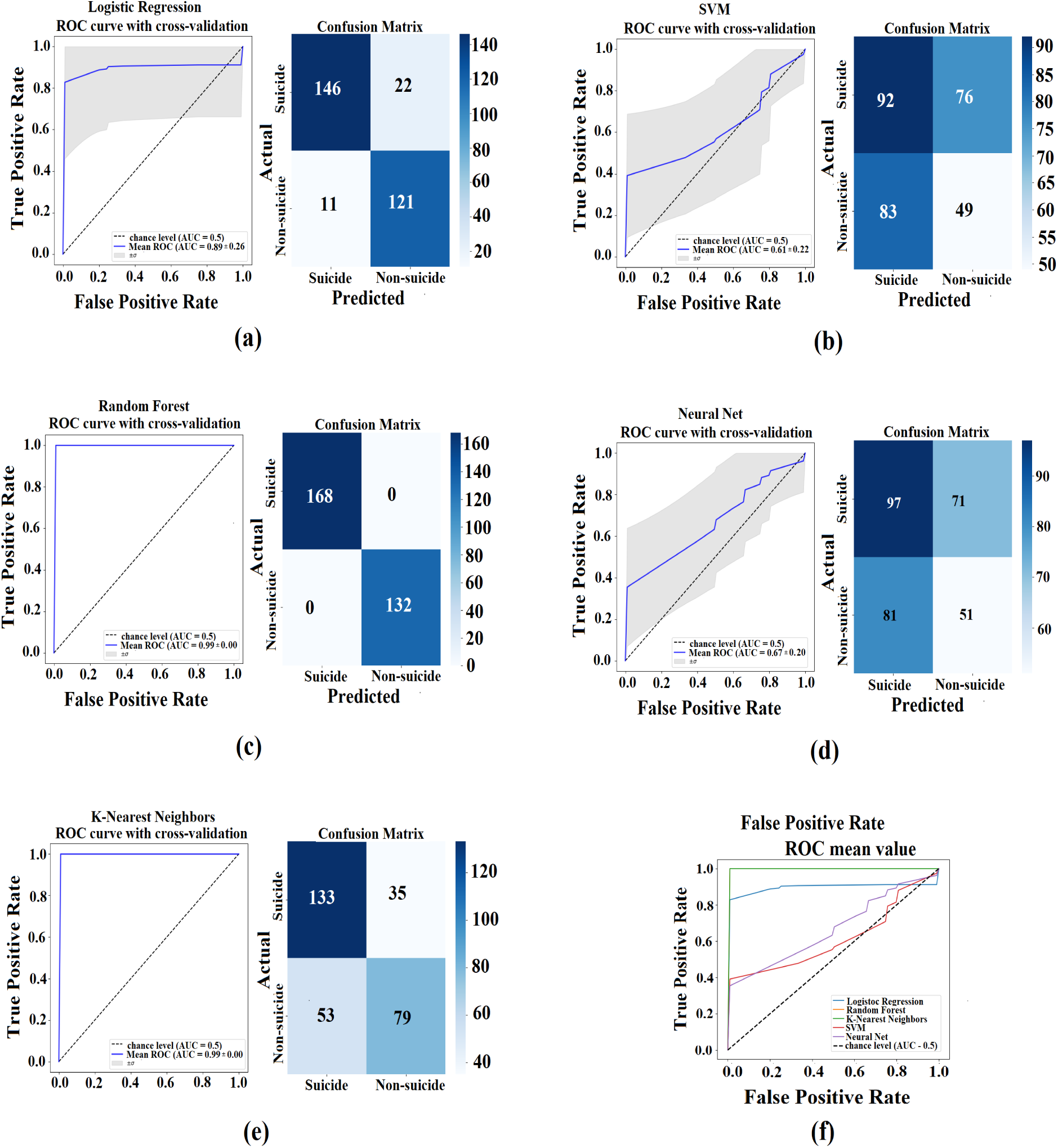
Prediction accuracy by varying the number of genes. (a) The variation in model accuracy for a logistic regression machine learning (ML) model in relation to the number of genes used as predictor features, ranging from 1 to 10 genes. (b) The accuracy of various ML models when specifically using 8 predictors.

The trained models were also used to assess the status of 7 BD patients whose conditions were not previously identified, yet some of their clinical data suggests that they are at a high risk of suicide, such as a first-degree relative who died of suicide and previous suicide attempts. For this classification, we took the majority decision by each of the classifiers (‘Combined Results’). The analysis revealed that patients ROU7776B and 21868 are at a low risk of suicide, whereas the other five BD patients are classified as at a high risk. These findings are detailed in Table 2. Interestingly, the BD patient with no previous suicide attempts and no family history was classified as a low risk for suicide. The other one that was classified as a low-risk had no previous attempts and one family member who died by suicide. The rest of the patients were classified as a high-risk for suicide. These patients had previously attempted suicide or had a few family members who died by suicide (with one patient 14501 having just one family member who died by suicide).

**Table 2:** Clinical Details of unknown labelled BD patients.

|  | No. of suicide attempts | Family history of Suicide | Logistic Regression | Random Forest | K-Nearest Neighbors | SVM | Neural Net | Combined Results |
| --- | --- | --- | --- | --- | --- | --- | --- | --- |
| R29425B | 2 | 1 | 1 | 1 | 1 | 1 | 1 | 1 |
| R34914A | 1 | 0 | 1 | 1 | 1 | 1 | 1 | 1 |
| Rou7776B | 0 | 0 | 0 | 0 | 0 | 1 | 1 | 0 |
| Rou6223B | 0 | 2 | 1 | 1 | 1 | 1 | 1 | 1 |
| R9288 | 0 | 3 | 1 | 1 | 1 | 1 | 1 | 1 |
| 14501 | 0 | 1 | 1 | 1 | 1 | 1 | 1 | 1 |
| 21868 | 0 | 1 | 0 | 0 | 0 | 1 | 0 | 0 |

## 4 Discussion

In this study, we have unearthed distinct gene expression signatures intricately linked to suicide and various facets of its phenomenology by utilizing RNA sequencing data from LCLs of BD patients who died by suicide compared to BD patients who are at a low risk of suicide. After identifying genes and pathways linked with suicide, we have further used these as features to train classification algorithms in order to construct a predictor for suicide risk in BD using biomarkers that are easily and cheaply available without much stress to the patient. We initially identified 843 genes that were differentially expressed in LCLs of BD patients who died by suicide compared to patients at a low risk of suicide, since they were followed and monitored over the years and have not attempted suicide or had suicidal thoughts. We used several pathways, disease, and functional annotation analyses to first identify possible mechanisms of suicide. Importantly, our biological samples were LCLs – immune-related cells. We therefore expected to find immune-related changes and did not expect to find brain-related implications. For brain-related mechanisms, we assumed that iPSC work related to differentiating the cells into neurons would be required. We were very surprised, therefore, to find many neuronal pathways that are altered between patients at a high risk of suicide and patients with the low risk. This suggests that when differentiating neurons, we will find neurophysiological changes in neurons derived from patients at a high risk of suicide compared to those at a low risk.

Among the primary findings, DEGs were enriched for the primary immunodeficiency pathway between LCL from BD patients who died by suicide and patients who are at a low risk of suicide. For example, genes such as *LCK* (*log*_2_ fold change of 0.81), *AICDA*, (*log*_2_ fold change of 1.329), and *CIITA* (*log*_2_ fold change of 0.975) exhibited increased expression levels in LCL from BD patients who died by suicide compared to those at a low risk of suicide. These genes are critical for immune system functioning; *LCK* is involved in T-cell receptor signaling [75]. *CIITA* (Class II Major Histocompatibility Complex Transactivator) regulates MHC (Major Histocompatibility complex) class II genes [76]. *AICDA* is crucial for B-cell antibody response [77]. The differential expression of these genes might reflect an underlying dysregulation in the immune response, which could be contributing to the pathophysiology of suicide. On the other hand, the *CD8A* gene (*log*_2_ fold change of 2.008), which is vital for T-cell mediated cytotoxicity [78], showed lower expression in LCLs from people who died by suicide, suggesting a potential protective role of this gene against suicide. This pattern of gene expression adds an intriguing layer to our understanding of how genes associated with both high and low risk can be linked to immune dysregulation and psychiatric conditions, particularly in the context of suicide in BD. Continued surveillance of these genes is crucial, as it will enable further exploration into the complex relationships between immune function disturbances and suicide.

Further supporting the link between immune function and the brain, our study also highlighted significant alterations in cell adhesion and ion channel pathways in LCLs of BD patients who died by suicide compared to the patients at a low risk. These pathways are known to influence neuronal connectivity and excitability [79, 80]. From the cell adhesion pathway, genes like *NTNG1* and *NRXN1* showed increased expression, with *log*_2_ fold changes of 2.27 and 1.42, respectively in LCLs of BD patients who died by suicide compared to BD patients with a low risk of suicide. Both genes are crucial for axonal guidance and synaptic stability [81, 82]. Additionally, *CD8A* was upregulated with a *log*_2_ fold change of 2.008 and. This gene also participates in the primary immunodeficiency pathway. The gene *CLDN10* is a key component of the blood-brain barrier (BBB) tight junctions [83] and was down-regulated with a *log*_2_ fold change of -2.16 in LCLs of people who died by suicide. Dysregulation in cell adhesion genes can lead to altered excitatory/inhibitory balance within the neural circuit [84–87].

Ion channels are crucial in modulating neuronal excitability and signal transduction, with growing evidence linking their dysregulation to psychiatric disorders [88, 89]. Our study highlights significant gene expression differences in BD patients who died by suicide compared to those with a low risk of suicide. Specifically, voltage-dependent calcium channel genes such as *CACNB2* (Calcium Voltage-Gated Channel Auxiliary Subunit Beta 2) showed a notable increase (*log*_2_ fold change of 1.318) in BD patients who died by suicide. In contrast, *CACNA1A* (Calcium Voltage-Gated Channel Sub-unit Alpha1 A) exhibited a *log*_2_ fold change of 0.955, predominantly in patients with a low risk of suicide. These genes mediate calcium influx into a wide variety of electrically excitable cells, including cardiac, muscle, neurons, and sensory cells [90]. In a similar pattern, *SCN11A* and *SCN1B* genes, which encode for voltage-gated sodium channels [91], showed differential expression. *SCN11A* (sodium voltage-gated channel alpha subunit 11) was up-regulated (*log*_2_ fold change of 1.355) in BD patients who died by suicide, while *SCN1B* (sodium voltage-gated channel beta subunit 1) showed a higher expression (*log*_2_ fold change of 2.33) in patients with a low risk of suicide. This differential expression of calcium and sodium channel genes suggests distinct roles, potentially categorizing them as either low or high predictors of suicide risk. Moreover, genes encoding for calcium-activated potassium channels such as *KCNN2* (Potassium Calcium-Activated Channel Subfamily N Member 2) and *KCNMB1* (Potassium Calcium-Activated Channel Subfamily M Regulatory Beta Subunit 1) were also highly expressed in BD patients who died by suicide [92] with a *log*_2_ fold change of 3.228 and 1.988 respectively. Additionally, the serotonin receptor gene *HTR3B* also showed a significant up-regulation (*log*_2_ fold change of 3.4) in BD patients who died by suicide, reaffirming the well-established link between serotonin signaling disruptions and both depressive and suicidal behaviors [42, 93–95]. These insights reveal a complex interplay between genetic factors and ion channel function, further elucidating the multifaceted nature of suicide risk in BD patients.

Likewise, our findings support the established link between ion channel dysfunctions and their role in both cardiovascular and neurological disorders, highlighting their critical roles in cellular excitability and signaling [96]. This connection has been well-documented, with ion channel abnormalities known to contribute to conditions such as cardiac arrhythmias and various neurological maladies, suggesting a shared pathophysiological foundation [97]. In our study, we observed that genes such as *KCNN2* (*log*_2_ fold change of 3.228), *KCNMB1* (*log*_2_ fold change of 1.98), *SCN11A* (*log*_2_ fold change of 1.35), *CACNB2* (*log*_2_ fold change of 1.318) and glutamate ionotropic receptor AMPA type subunit 1 also known as *GRIA1* (*log*_2_ fold change of 2.85), showed increased expression in LCLs from BD patients who died by suicide, while a few ion channels like *SCN1B* (*log*_2_ fold change of 2.33) and *KCNE4* (*log*_2_ fold change of 4.06) were more prominently expressed in BD patients with a low risk of suicide. This pattern of differential gene expression underscores the significant role these channels play in influencing both neuronal and cardiovascular functions, which in turn affect mood and behavior. The data points to a complex interaction between genetic factors that impact both neurological health and cardiovascular stability, reinforcing the potential of these genes as dual biomarkers for assessing the risk of suicide as well as cardiovascular diseases.

We also identified genes associated with astrocytosis and gliosis. Astrocytosis and gliosis involve the proliferation and hypertrophy of astrocytes, usually in response to central nervous system pathologies like neurodegenerative disease, tumor growth, and trauma [98]. We found five genes to be differentially expressed, among which *BDNF* and *ITGA5* were notably upregulated in BD patients who are at a low risk of suicide, with *log*_2_ fold changes of 2.04 and 1.30, respectively. *BDNF* is pivotal for neuronal survival and development, influencing the growth, differentiation, and synaptic plasticity of neurons [99]. It also plays a critical role in neuronal and synaptic regeneration and repair following injury [100]. On the other hand, *ITGA5* is essential for cell adhesion to the extracellular matrix, supporting cell migration during development and wound healing, and is involved in angiogenesis [101, 102]. The roles of neurotrophins like *BDNF* and integrins like *ITGA5* are crucial in guiding neuron positioning, growth, and maturation, highlighting why their low expression might be observed in suicide samples [103, 104]. However, the *GRM8* gene was found to be elevated in LCLS of BD patients who died by suicide (*log*_2_ fold change of 1.38), and encodes for glutamate metabotropic receptor 8 and plays a significant role in glutamate neurotransmission. This gene’s function is particularly pertinent in BD, where glutamate system dysregulation is often implicated [105]. Additionally, the *GRM8* gene’s involvement in processes like astrocytosis can be linked to altered glutamatergic signaling which may influence the proliferation and hypertrophy of astrocytes during neuroinflammatory responses.

Intriguingly, beyond their hematopoietic lineage, LCLs exhibit a remarkable expression profile reminiscent of neuronal activity, as evidenced by RNA sequencing studies [106, 107]. This unexpected convergence underscores the potential of LCLs as a valuable model for probing the genetic underpinnings of neuropsychiatric and neurodevelopmental disorders [108–110]. In our analysis, several genes, including *SHANK2* (*log*_2_ fold change 1.96), *CACNG5* (*log*_2_ fold change 4.93), *ANK3* (*log*_2_ fold change 2.04), and *NRG3*, (*log*_2_ fold change 4.29) were found to be significantly upregulated in LCLs of BD patients who died by suicide compared to patients who are at a low risk of suicide. *SHANK2* plays a crucial role in synaptic transmission and has been associated with autism spectrum disorders and schizophrenia [111, 112]. *CACNG5* encodes a subunit of the voltage-dependent calcium channels [113] and has been previously known to be linked to schizophrenia [114]. *ANK3* is crucial for encoding ankyrin-G, a protein involved in the structural stability of neurons by anchoring integral membrane proteins to the cytoskeleton [115]. Previous studies have shown that variations in this gene are associated with both BD and schizophrenia [116]. *NRG3*, encoding neuregulin 3, is a part of the neuregulin family. Neuregulins act through binding to the ERBB family initiating important signalling pathways that influence cell growth, and differentiation [117]. Genetic studies have linked variations in the *NRG3* gene to schizophrenia and BD [118]. This concurrent association underscores the potential role of these genes as high-risk factors for suicide in BD. The upregulation of these genes not only reinforces their critical function in neuropsychiatric pathologies but also highlights them as promising biomarkers for assessing suicide risk.

To delve deeper, we enhanced our understanding of the genetic factors that influence BD, particularly in distinguishing between ‘SUICIDE’ and ‘NON-SUICIDE’ tendencies, by refining our research methodology through a robust protocol. Out of 843 DEGs, we selected 50 potential biomarkers to be analyzed via machine learning techniques. We established a list of the top 10 markers, which showcased consistently high predictive accuracy. These genes are *LYL1*, *LMO7*, *DDAH2*, *PAG1*, *LCK*, *AP001372.2*, *INPP5D*, *SNX8*, *TBC1D4*, and *GJC1*. Importantly, these top 10 genes differ from the initial 843 DEGs, showing clearer gene count distinctions between ‘SUICIDE’ and ‘NON-SUICIDE’ cases. Our methodology involved evaluating the performance of machine learning models ranging from 1 to 10 predictors to validate the robustness of these biomarkers.

Our findings were particularly significant in the evaluation of predictive performance across various machine learning models. When narrowed down to an optimized set of eight specific genes from our top 10, we observed enhanced prediction accuracies. The accuracy levels achieved by different models were as follows: Logistic Regression at 98.3%*±*5.0%, Random Forest at 96.0%*±*10.8%, K-Nearest Neighbors at 77.0%*±*30.7%, Support Vector Machine at 90.7%*±*20.6%, and Neural Networks at an impressive 99.7%*±*2.3%. These findings emphasize both the robustness and reliability of our chosen genetic markers and underscore their clinical effectiveness in distinguishing between ‘SUICIDE’ and ‘NON-SUICIDE’ BD patients. The trained models were used finally to assess 7 BD patients, previously un-diagnosed but identified as high-risk due to factors like family history of suicide and personal attempts. Using a majority decision from the classifiers, the analysis indicated that two patients, ROU7776B and 21868, are at a low risk of suicide, while the remaining five are at high risk. Interestingly, Rou7776B has never attempted suicide, nor did he have any family history of suicide. Patient 21868 attempted suicide once and had one family member who died by suicide. Most high-risk patients either had multiple personal attempts or multiple family members with a history of suicide (except for patient 14501, who had one only one family member). This implies that our findings could contribute to the development of targeted therapeutic strategies and improve preventive measures in BD management. These aspects are crucial for enhancing patient outcomes and could potentially lead to personalized treatment approaches based on genetic predispositions.

## 5 Conclusion

In conclusion, our study provides valuable insights into the genetic underpinnings of suicide in BD, elucidating complex interactions among genes involved in primary immunodeficiency, cellular signalling pathways such as cell adhesion and ion channels, and their links to other conditions like cardiovascular disorders. Through rigorous analysis of RNA sequencing data, we identified distinct gene expression signatures associated with low and high risks of suicide in BD patients and constructed high accuracy classifiers, highlighting potential biomarkers that could lead to more precise, accessible, and stress-free diagnostic methods. This approach underscores the potential of integrating genetic and computational methods to advance our understanding and management of psychiatric disorders, aiming toward personalized therapeutic strategies to improve patient outcomes.

## Supporting information

https://github.com/omveersharmanet/Predicting-Suicide-Risk-in-Bipolar-Disorder-patients.git

## Acknowledgements

The graphical image for this publication was created with Biorender.com and the figures were plotted in MATLAB.

## Declarations

### Funding

The Zuckerman STEM leadership program and Israel Science Foundation grants - 1994/21 and 3252 /21 for Prof. Shani Stern.

### Conflict of interest/Competing interests

The authors declare no competing interests

### Ethics approval and consent to participate

All participants provided informed consent before participating in the study. Approval for the study was obtained from the Research Ethics Board of the University of Cagliari, Italy.

### Author contribution

Omveer Sharma (OS) and Ritu Nayak (RN) (co-first authors) contributed equally to this work. OS performed the data analysis, conducted machine learning predictor analysis, and drafted and compiled the manuscript. RN grew the cells, performed RNA extraction experiments, and drafted and compiled the manuscript. Wote Amelo Rike (WAR) and Ashwani Choudhary (AC) assisted with cell culture. Yara Hussein (YH) and Aviram Shemen (AS) assisted with data analysis. Idan Rosh (IR) provided cell care. Utkarsh Tripathi (UT) reviewed the manuscript. Alessio Squassina (AS) provided the clinical samples for the cohort study. Martin Alda (MA) contributed to the study design and provided resources. Shani Stern (SS) (corresponding author) conceptualized and supervised the study, analyzed the data, and drafted the manuscript.

All authors reviewed the manuscript.

### Data availability

Correspondence and requests for materials should be addressed to Prof. Shani Stern

### Materials availability

Correspondence and requests for materials should be addressed to Prof. Shani Stern

### Code availability

Correspondence and requests for materials should be addressed to Prof. Shani Stern. Additional information about the samples will be available in the GitHub file at this link (https://github.com/omveersharmanet/Predicting-Suicide-Risk-in-Bipolar-Disorder-patients.git).

## Notes

https://github.com/omveersharmanet/Predicting-Suicide-Risk-in-Bipolar-Disorder-patients.git

### Competing Interest Statement

The authors have declared no competing interest.

## References

1. Suicide.: https://www.who.int/news-room/fact-sheets/detail/suicide

[2] Bachmann, S.: Epidemiology of suicide and the psychiatric perspective. International journal of environmental research and public health 15(7), 1425 (2018)

[3] Pinals, D.A.: Liability and patient suicide. Focus 17(4), 349–354 (2019)

[4] Cavanagh, J.T., Carson, A.J., Sharpe, M., Lawrie, S.M.: Psychological autopsy studies of suicide: a systematic review. Psychological medicine 33(3), 395–405 (2003)

[5] Arsenault-Lapierre, G., Kim, C., Turecki, G.: Psychiatric diagnoses in 3275 suicides: a meta-analysis. BMC psychiatry 4, 1–11 (2004)

[6] Harris, E.C., Barraclough, B.: Suicide as an outcome for mental disorders. a meta-analysis. British journal of psychiatry 170(3), 205–228 (1997)

[7] Jones, S., Riste, L., Barrowclough, C., Bartlett, P., Clements, C., Davies, L., Holland, F., Kapur, N., Lobban, F., Long, R., et al.: Reducing relapse and suicide in bipolar disorder: practical clinical approaches to identifying risk, reducing harm and engaging service users in planning and delivery of care–the parades (psychoeducation, anxiety, relapse, advance directive evaluation and suicidality) programme (2018)

[8] Silva Costa, L., Alencar, A.P., Neto, P.J.N., Santos, M.d.S.V., Silva, C.G.L., Pinheiro, S.d.F.L., Silveira, R.T., Bianco, B.A.V., Júnior, R.F.F.P., Lima, M.A.P., et al.: Risk factors for suicide in bipolar disorder: a systematic review. Journal of affective disorders 170, 237–254 (2015)

[9] Eroglu, M.Z., Karakus, G., Tamam, L.: Bipolar disorder and suicide. Dusunen Adam 26(2), 139 (2013)

[10] Simon, N.M., Zalta, A.K., Otto, M.W., Ostacher, M.J., Fischmann, D., Chow, C.W., Thompson, E.H., Stevens, J.C., Demopulos, C.M., Nierenberg, A.A., et al.: The association of comorbid anxiety disorders with suicide attempts and suicidal ideation in outpatients with bipolar disorder. Journal of psychiatric research 41(3-4), 255–264 (2007)

[11] Plans, L., Barrot, C., Nieto, E., Rios, J., Schulze, T., Papiol, S., Mitjans, M., Vieta, E., Benabarre, A.: Association between completed suicide and bipolar disorder: A systematic review of the literature. Journal of affective disorders 242, 111–122 (2019)

[12] Zhu, R., Tian, S., Wang, H., Jiang, H., Wang, X., Shao, J., Wang, Q., Yan, R., Tao, S., Liu, H., et al.: Discriminating suicide attempters and predicting suicide risk using altered frontolimbic resting-state functional connectivity in patients with bipolar ii disorder. Frontiers in psychiatry 11, 597770 (2020)

[13] Strakowski, S.M., McElroy, S.L., Keck Jr, P.E., West, S.A.: Suicidality among patients with mixed and manic bipolar disorder. The American journal of psychiatry 153(5), 674–676 (1996)

[14] Dome, P., Rihmer, Z., Gonda, X.: Suicide risk in bipolar disorder: a brief review. Medicina 55(8), 403 (2019)

[15] Lee, S., Tsang, A., Kessler, R.C., Jin, R., Sampson, N., Andrade, L., Karam, E.G., Mora, M.E.M., Merikangas, K., Nakane, Y., et al.: Rapid-cycling bipolar disorder: cross-national community study. The British Journal of Psychiatry 196(3), 217–225 (2010)

[16] Berardelli, I., Rogante, E., Sarubbi, S., Erbuto, D., Cifrodelli, M., Concolato, C., Pasquini, M., Lester, D., Innamorati, M., Pompili, M.: Is lethality different between males and females? clinical and gender differences in inpatient suicide attempters. International journal of environmental research and public health 19(20), 13309 (2022)

[17] Monnin, J., Thiemard, E., Vandel, P., Nicolier, M., Tio, G., Courtet, P., Bellivier, F., Sechter, D., Haffen, E.: Sociodemographic and psychopathological risk factors in repeated suicide attempts: gender differences in a prospective study. Journal of affective disorders 136(1-2), 35–43 (2012)

[18] Miller, J.N., Black, D.W.: Bipolar disorder and suicide: a review. Current psychiatry reports 22, 1–10 (2020)

[19] Isometsä, E.T., Lönnqvist, J.K.: Suicide attempts preceding completed suicide. The British Journal of Psychiatry 173(6), 531–535 (1998)

[20] Bostwick, J.M., Pankratz, V.S.: Affective disorders and suicide risk: a reexamination. American Journal of Psychiatry 157(12), 1925–1932 (2000)

[21] Conwell, Y., Duberstein, P.R., Cox, C., Herrmann, J., Forbes, N., Caine, E.D.: Age differences in behaviors leading to completed suicide. The American Journal of Geriatric Psychiatry 6(2), 122–126 (1998)

[22] Ribeiro, J.D., Franklin, J.C., Fox, K.R., Bentley, K.H., Kleiman, E.M., Chang, B.P., Nock, M.K.: Self-injurious thoughts and behaviors as risk factors for future suicide ideation, attempts, and death: a meta-analysis of longitudinal studies. Psychological medicine 46(2), 225–236 (2016)

[23] Antypa, N., Serretti, A.: Family history of a mood disorder indicates a more severe bipolar disorder. Journal of affective disorders 156, 178–186 (2014)

[24] Antypa, N., Antonioli, M., Serretti, A.: Clinical, psychological and environmental predictors of prospective suicide events in patients with bipolar disorder. Journal of psychiatric research 47(11), 1800–1808 (2013)

[25] Guillaume, S., Jaussent, I., Jollant, F., Rihmer, Z., Malafosse, A., Courtet, P.: Suicide attempt characteristics may orientate toward a bipolar disorder in attempters with recurrent depression. Journal of affective disorders 122(1-2), 53–59 (2010)

[26] Erten, E., Uney, A.F., Saatçioğlu, Ö., Ozdemir, A., Fıstıkçı, N., Çakmak, D.: Effects of childhood trauma and clinical features on determining quality of life in patients with bipolar i disorder. Journal of affective disorders 162, 107–113 (2014)

[27] Maniglio, R.: The impact of child sexual abuse on the course of bipolar disorder: a systematic review. Bipolar disorders 15(4), 341–358 (2013)

[28] Etain, B., Aas, M., Andreassen, O.A., Lorentzen, S., Dieset, I., Gard, S., Kahn, J.-P., Bellivier, F., Leboyer, M., Melle, I., et al.: Childhood trauma is associated with severe clinical characteristics of bipolar disorders. The Journal of clinical psychiatry 74(10), 2585 (2013)

[29] Aas, M., Etain, B., Bellivier, F., Henry, C., Lagerberg, T., Ringen, A., Agartz, I., Gard, S., Kahn, J.-P., Leboyer, M., et al.: Additive effects of childhood abuse and cannabis abuse on clinical expressions of bipolar disorders. Psychological medicine 44(8), 1653–1662 (2014)

[30] Tsai, S.-Y., Lee, J.-C., Chen, C.-C.: Characteristics and psychosocial problems of patients with bipolar disorder at high risk for suicide attempt. Journal of Affective Disorders 52(1-3), 145–152 (1999)

[31] Chiba, T., Ide, K., Murakami, M., Kobayashi, N., Oka, T., Nakai, F., Yorizawa, R., Miyake, Y., Hamamura, T., Honjo, M., et al.: Event-related ptsd symptoms as a high-risk factor for suicide: longitudinal observational study. Nature Mental Health 1(12), 1013–1022 (2023)

[32] Nayak, R., Rosh, I., Kustanovich, I., Stern, S.: Mood stabilizers in psychiatric disorders and mechanisms learnt from in vitro model systems. International journal of molecular sciences 22(17), 9315 (2021)

[33] Tobe, B.T., Crain, A.M., Winquist, A.M., Calabrese, B., Makihara, H., Zhao, W.-n., Lalonde, J., Nakamura, H., Konopaske, G., Sidor, M., et al.: Probing the lithium-response pathway in hipscs implicates the phosphoregulatory set-point for a cytoskeletal modulator in bipolar pathogenesis. Proceedings of the National Academy of Sciences 114(22), 4462–4471 (2017)

[34] Stern, S., Santos, R., Marchetto, M., Mendes, A., Rouleau, G., Biesmans, S., Wang, Q., Yao, J., Charnay, P., Bang, A., et al.: Neurons derived from patients with bipolar disorder divide into intrinsically different sub-populations of neurons, predicting the patients’ responsiveness to lithium. Molecular psychiatry 23(6), 1453–1465 (2018)

[35] Stern, S., Sarkar, A., Stern, T., Mei, A., Mendes, A.P.D., Stern, Y., Goldberg, G., Galor, D., Nguyen, T., Randolph-Moore, L., Kim, Y., Rouleau, G., Bang, A., Alda, M., Santos, R., Marchetto, M.C., Gage, F.H.: Mechanisms underlying the hyperexcitability of ca3 and dentate gyrus hippocampal neurons derived from patients with bipolar disorder. Biological Psychiatry 88(2), 139–149 (2020) 10.1016/j.biopsych.2019.09.018. Mechanisms of Mood Disorders

[36] Stern, S., Sarkar, A., Galor, D., Stern, T., Mei, A., Stern, Y., Mendes, A.P., Randolph-Moore, L., Rouleau, G., Bang, A.G., et al.: A physiological instability displayed in hippocampal neurons derived from lithium-nonresponsive bipolar disorder patients. Biological psychiatry 88(2), 150–158 (2020)

[37] Tripathi, U., Mizrahi, L., Alda, M., Falkovich, G., Stern, S.: Information theory characteristics improve the prediction of lithium response in bipolar disorder patients using a support vector machine classifier. Bipolar Disorders 25(2), 110–127 (2023)

[38] Mizrahi, L., Choudhary, A., Ofer, P., Goldberg, G., Milanesi, E., Kelsoe, J.R., Gurwitz, D., Alda, M., Gage, F.H., Stern, S.: Immunoglobulin genes expressed in lymphoblastoid cell lines discern and predict lithium response in bipolar disorder patients. Molecular Psychiatry, 1–14 (2023)

[39] Nayak, R., Rosh, I., Rabinski, T., Falik, D., Percia, M.M., Stern, S.: Generation and characterization of ipsc lines (uohi003-a, uohi002-a) from a patient with shank3 mutation and her healthy mother. Stem Cell Research 64, 102899 (2022)

[40] Hatoum, A.S., Colbert, S.M., Johnson, E.C., Huggett, S.B., Deak, J.D., Pathak, G.A., Jennings, M.V., Paul, S.E., Karcher, N.R., Hansen, I., et al.: Multivariate genome-wide association meta-analysis of over 1 million subjects identifies loci underlying multiple substance use disorders. Nature mental health 1(3), 210–223 (2023)

[41] Choudhary, A., Peles, D., Nayak, R., Mizrahi, L., Stern, S.: Current progress in understanding schizophrenia using genomics and pluripotent stem cells: A meta-analytical overview. Schizophrenia Research (2022)

[42] Antypa, N., Serretti, A., Rujescu, D.: Serotonergic genes and suicide: A systematic review. European Neuropsychopharmacology 23(10), 1125–1142 (2013) 10.1016/j.euroneuro.2013.03.013

[43] Brezo, J., Klempan, T., Turecki, G.: The genetics of suicide: A critical review of molecular studies. Psychiatric Clinics of North America 31(2), 179–203 (2008) 10.1016/j.psc.2008.01.008. Suicidal Behavior: A Developmental Perspective

[44] Courtet, P., Jollant, F., Castelnau, D., Buresi, C., Malafosse, A.: Suicidal behavior: relationship between phenotype and serotonergic genotype. In: American Journal of Medical Genetics Part C: Seminars in Medical Genetics, vol. 133, pp. 25–33 (2005). Wiley Online Library

[45] Brezo, J., Bureau, A., Mérette, C., Jomphe, V., Barker, E.D., Vitaro, F., Hébert, M., Carbonneau, R., Tremblay, R.E., Turecki, G.: Differences and similarities in the serotonergic diathesis for suicide attempts and mood disorders: a 22-year longitudinal gene–environment study. Molecular psychiatry 15(8), 831–843 (2010)

[46] Mann, J.J.: The serotonergic system in mood disorders and suicidal behaviour. Philosophical Transactions of the Royal Society B: Biological Sciences 368(1615), 20120537 (2013)

[47] Dwivedi, Y.: Brain-derived neurotrophic factor and suicide pathogenesis. Annals of medicine 42(2), 87–96 (2010)

[48] Banerjee, R., Ghosh, A.K., Ghosh, B., Bhattacharyya, S., Mondal, A.C.: Decreased mrna and protein expression of bdnf, ngf, and their receptors in the hippocampus from suicide: an analysis in human postmortem brain. Clinical Medicine Insights: Pathology 6, 12530 (2013)

[49] Kohli, M.A., Salyakina, D., Pfennig, A., Lucae, S., Horstmann, S., Menke, A., Kloiber, S., Hennings, J., Bradley, B.B., Ressler, K.J., et al.: Association of genetic variants in the neurotrophic receptor–encoding gene ntrk2 and a lifetime history of suicide attempts in depressed patients. Archives of general psychiatry 67(4), 348–359 (2010)

[50] Pivac, N., Pregelj, P., Nikolac, M., Zupanc, T., Nedic, G., Muck Seler, D., Videtic Paska, A.: The association between catechol-o-methyl-transferase val108/158met polymorphism and suicide. Genes, brain and behavior 10(5), 565–569 (2011)

[51] Antypa, N., Souery, D., Tomasini, M., Albani, D., Fusco, F., Mendlewicz, J., Serretti, A.: Clinical and genetic factors associated with suicide in mood disorder patients. European archives of psychiatry and clinical neuroscience 266, 181–193 (2016)

[52] Isayeva, U., Manchia, M., Collu, R., Primavera, D., Deriu, L., Caboni, E., Iaselli, N., Sundas, D., Tusconi, M., Pinna, F., et al.: Exploring the association between brain-derived neurotrophic factor levels and longitudinal psychopathological and cognitive changes in sardinian psychotic patients. European Psychiatry 65(1), 71 (2022)

[53] Magno, L., Miranda, D., Neves, F., Pimenta, G., Mello, M., De Marco, L., Correa, H., Romano-Silva, M.A.: Association between akt1 but not aktip genetic variants and increased risk for suicidal behavior in bipolar patients. Genes, brain and behavior 9(4), 411–418 (2010)

[54] Ekinci, O., Albayrak, Y., Ekinci, A.E., Caykoylu, A.: Relationship of trait impulsivity with clinical presentation in euthymic bipolar disorder patients. Psychiatry research 190(2-3), 259–264 (2011)

[55] Jiménez, E., Arias, B., Mitjans, M., Goikolea, J., Roda, E., Saiz, P., García-Portilla, M., Burón, P., Bobes, J., Oquendo, M., et al.: Genetic variability at impa2, inpp1 and gsk3*β* increases the risk of suicidal behavior in bipolar patients. European Neuropsychopharmacology 23(11), 1452–1462 (2013)

[56] Sequeira, A., Mamdani, F., Lalovic, A., Anguelova, M., Lesage, A., Seguin, M., Chawky, N., Desautels, A., Turecki, G.: Alpha 2a adrenergic receptor gene and suicide. Psychiatry research 125(2), 87–93 (2004)

[57] Zai, C.C., Gonçalves, V.F., Tiwari, A.K., Gagliano, S.A., Hosang, G., De Luca, V., Shaikh, S.A., King, N., Chen, Q., Xu, W., et al.: A genome-wide association study of suicide severity scores in bipolar disorder. Journal of psychiatric research 65, 23–29 (2015)

[58] Docherty, A.R., Mullins, N., Ashley-Koch, A.E., Qin, X., Coleman, J.R., Shabalin, A., Kang, J., Murnyak, B., Wendt, F., Adams, M., et al.: Gwas meta-analysis of suicide attempt: identification of 12 genome-wide significant loci and implication of genetic risks for specific health factors. American journal of psychiatry 180(10), 723–738 (2023)

[59] Darlington, G.J.: Epstein-barr virus transformation of lymphoblasts. CSH Protoc 2006, 4481 (2006)

[60] Andrews, S., et al.: FastQC: a quality control tool for high throughput sequence data. Cambridge, United Kingdom (2010)

[61] Dobin, A., Davis, C.A., Schlesinger, F., Drenkow, J., Zaleski, C., Jha, S., Batut, P., Chaisson, M., Gingeras, T.R.: Star: ultrafast universal rna-seq aligner. Bioinformatics 29(1), 15–21 (2013)

[62] Love, M.I., Huber, W., Anders, S.: Moderated estimation of fold change and dispersion for rna-seq data with deseq2. Genome biology 15, 1–21 (2014)

[63] Benjamini, Y., Hochberg, Y.: Controlling the false discovery rate: a practical and powerful approach to multiple testing. Journal of the Royal statistical society: series B (Methodological) 57(1), 289–300 (1995)

[64] Gao, M., Wong, N.M., Lin, C., Huang, C.-M., Liu, H.-L., Toh, C.-H., Wu, C., Tsai, Y.-F., Lee, S.-H., Lee, T.M.: Multimodal brain connectome-based prediction of suicide risk in people with late-life depression. Nature mental health 1(2), 100–113 (2023)

[65] Sharma, O., Sahoo, N.C., Puhan, N.B.: Highway lane-changing prediction using a hierarchical software architecture based on support vector machine and continuous hidden markov model. International Journal of Intelligent Transportation Systems Research 20(2), 519–539 (2022)

[66] Lalvani, S., Bari, S., Vike, N.L., Stefanopoulos, L., Kim, B.-W., Block, M., Maglaveras, N., Katsaggelos, A.K., Breiter, H.C.: Predicting suicidality with small sets of interpretable reward behavior and survey variables. Nature Mental Health, 1–14 (2024)

[67] Madanlal, D., Guinard, C., Nuñez, V.P., Becker, S., Garnham, J., Khayachi, A., Léger, S., O’Donovan, C., Singh, S., Stern, S., et al.: A pilot study examining the impact of lithium treatment and responsiveness on mnemonic discrimination in bipolar disorder. Journal of Affective Disorders 351, 49–57 (2024)

[68] Singh, S., Khayachi, A., Stern, S., Trappenberg, T., Alda, M., Nunes, A.: The effects of bipolar disorder granule cell hyperexcitability and lithium therapy on pattern separation in a computational model of the dentate gyrus. bioRxiv, 2024–04 (2024)

[69] Santos, R., Linker, S.B., Stern, S., Mendes, A.P., Shokhirev, M.N., Erikson, G., Randolph-Moore, L., Racha, V., Kim, Y., Kelsoe, J.R., et al.: Deficient lef1 expression is associated with lithium resistance and hyperexcitability in neurons derived from bipolar disorder patients. Molecular psychiatry 26(6), 2440–2456 (2021)

[70] Trivedi, D., Sharma, O., Pattnaik, S., Hazra, V., Puhan, N.B.: Improving rainfall forecast at the district scale over the eastern indian region using deep neural network. Theoretical and Applied Climatology 155(1), 761–777 (2024)

[71] Jeremian, R., Malinowski, A., Chaudhary, Z., Srivastava, A., Qian, J., Zai, C., Adanty, C., Fischer, C.E., Burhan, A.M., Kennedy, J.L., et al.: Epigenetic age dysregulation in individuals with bipolar disorder and schizophrenia. Psychiatry Research 315, 114689 (2022)

[72] Herrera-Rivero, M., Gutiérrez-Fragoso, K., Kurtz, J., Baune, B.T.: Immunogenetics of lithium response and psychiatric phenotypes in patients with bipolar disorder. Translational psychiatry 14(1), 174 (2024)

[73] Nielsen, R.E., Banner, J., Jensen, S.E.: Cardiovascular disease in patients with severe mental illness. Nature Reviews Cardiology 18(2), 136–145 (2021)

[74] Quraishi, I.H., Stern, S., Mangan, K.P., Zhang, Y., Ali, S.R., Mercier, M.R., Marchetto, M.C., McLachlan, M.J., Jones, E.M., Gage, F.H., et al.: An epilepsy-associated kcnt1 mutation enhances excitability of human ipsc-derived neurons by increasing slack kna currents. Journal of Neuroscience 39(37), 7438–7449 (2019)

[75] Shah, K., Al-Haidari, A., Sun, J., Kazi, J.U.: T cell receptor (tcr) signaling in health and disease. Signal transduction and targeted therapy 6(1), 412 (2021)

[76] Ting, J.P.-Y., Trowsdale, J.: Genetic control of mhc class ii expression. Cell 109(2), 21–33 (2002)

[77] Park, S.-R.: Activation-induced cytidine deaminase in b cell immunity and cancers. Immune Network 12(6), 230–239 (2012)

[78] Koh, C.-H., Lee, S., Kwak, M., Kim, B.-S., Chung, Y.: Cd8 t-cell subsets: heterogeneity, functions, and therapeutic potential. Experimental & Molecular Medicine 55(11), 2287–2299 (2023)

[79] Sheng, L., Leshchyns’ ka, I., Sytnyk, V.: Cell adhesion and intracellular calcium signaling in neurons. Cell Communication and Signaling 11, 1–13 (2013)

[80] Brown, D.A.: Neurons, receptors, and channels. Annual Review of Pharmacology and Toxicology 60, 9–30 (2020)

[81] Dean, C., Scholl, F.G., Choih, J., DeMaria, S., Berger, J., Isacoff, E., Scheiffele, P.: Neurexin mediates the assembly of presynaptic terminals. Nature neuroscience 6(7), 708–716 (2003)

[82] Zheng, M., Chen, R., Chen, H., Zhang, Y., Chen, J., Lin, P., Lan, Q., Yuan, Q., Lai, Y., Jiang, X., et al.: Netrin-1 promotes synaptic formation and axonal regeneration via jnk1/c-jun pathway after the middle cerebral artery occlusion. Frontiers in Cellular Neuroscience 12, 13 (2018)

[83] Berselli, A., Benfenati, F., Maragliano, L., Alberini, G.: Multiscale modelling of claudin-based assemblies: a magnifying glass for novel structures of biological interfaces. Computational and Structural Biotechnology Journal 20, 5984–6010 (2022)

[84] Kim, H.-G., Kishikawa, S., Higgins, A.W., Seong, I.-S., Donovan, D.J., Shen, Y., Lally, E., Weiss, L.A., Najm, J., Kutsche, K., et al.: Disruption of neurexin 1 associated with autism spectrum disorder. The American Journal of Human Genetics 82(1), 199–207 (2008)

[85] Gregor, A., Albrecht, B., Bader, I., Bijlsma, E.K., Ekici, A.B., Engels, H., Hackmann, K., Horn, D., Hoyer, J., Klapecki, J., et al.: Expanding the clinical spectrum associated with defects in cntnap2 and nrxn1. BMC medical genetics 12, 1–12 (2011)

[86] Chen, J., Dong, B., Feng, X., Jiang, D., Chen, G., Long, C., Yang, L.: Aberrant mpfc gabaergic synaptic transmission and fear behavior in neuroligin-2 r215h knock-in mice. Brain research 1730, 146671 (2020)

[87] Jesudas, B.R., Nandeesha, H., Menon, V., Allimuthu, P.: Relationship of elevated neural cell adhesion molecule 1 with interleukin-10 and disease severity in bipolar disorder. Asian Journal of Psychiatry 47, 101849 (2020)

[88] Dai, G.: Signaling by ion channels: Pathways, dynamics and channelopathies. Missouri Medicine 120(5), 367 (2023)

[89] Stern, S., Segal, M., Moses, E.: Involvement of potassium and cation channels in hippocampal abnormalities of embryonic ts65dn and tc1 trisomic mice. EBioMedicine 2(9), 1048–1062 (2015)

[90] Catterall, W.A.: Voltage-gated calcium channels. Cold Spring Harbor perspectives in biology 3(8), 003947 (2011)

[91] Bouza, A.A., Isom, L.L.: Voltage-gated sodium channel β subunits and their related diseases. Voltage-gated Sodium Channels: Structure, Function and Channelopathies, 423–450 (2018)

[92] Kim, D.M., Nimigean, C.M.: Voltage-gated potassium channels: a structural examination of selectivity and gating. Cold Spring Harbor perspectives in biology 8(5), 029231 (2016)

[93] Frank, B., Niesler, B., Nöthen, M.M., Neidt, H., Propping, P., Bondy, B., Rietschel, M., Maier, W., Albus, M., Rappold, G.: Investigation of the human serotonin receptor gene htr3b in bipolar affective and schizophrenic patients. American Journal of Medical Genetics Part B: Neuropsychiatric Genetics 131(1), 1–5 (2004)

[94] Wang, L., Wang, M., Zhao, C., Jian, J., Qiao, D.: Association of htr3b gene polymorphisms with depression and its executive dysfunction: a case–control study. BMC psychiatry 23(1), 128 (2023)

[95] Vadodaria, K.C., Stern, S., Marchetto, M.C., Gage, F.H.: Serotonin in psychiatry: in vitro disease modeling using patient-derived neurons. Cell and tissue research 371, 161–170 (2018)

[96] Kim, J.-B.: Channelopathies. Korean journal of pediatrics 57(1), 1 (2014)

[97] Bernardi, J., Aromolaran, K.A., Aromolaran, A.S.: Neurological disorders and risk of arrhythmia. International Journal of Molecular Sciences 22(1), 188 (2020)

[98] Pekny, M., Nilsson, M.: Astrocyte activation and reactive gliosis. Glia 50(4), 427–434 (2005)

[99] Park, H., Poo, M.-m.: Neurotrophin regulation of neural circuit development and function. Nature Reviews Neuroscience 14(1), 7–23 (2013)

[100] Nagappan, P.G., Chen, H., Wang, D.-Y.: Neuroregeneration and plasticity: a review of the physiological mechanisms for achieving functional recovery postinjury. Military Medical Research 7, 1–16 (2020)

[101] Sun, Z., Costell, M., Fässler, R.: Integrin activation by talin, kindlin and mechanical forces. Nature cell biology 21(1), 25–31 (2019)

[102] Plow, E.F., Meller, J., Byzova, T.V.: Integrin function in vascular biology: a view from 2013. Current opinion in hematology 21(3), 241–247 (2014)

[103] Wang, C.S., Kavalali, E.T., Monteggia, L.M.: Bdnf signaling in context: From synaptic regulation to psychiatric disorders. Cell 185(1), 62–76 (2022)

[104] Zou, Y., Zhang, Y., Tu, M., Ye, Y., Li, M., Ran, R., Zou, Z.: Brain-derived neurotrophic factor levels across psychiatric disorders: A systemic review and network meta-analysis. Progress in Neuro-Psychopharmacology and Biological Psychiatry, 110954 (2024)

[105] Blacker, C.J., Lewis, C.P., Frye, M.A., Veldic, M.: Metabotropic glutamate receptors as emerging research targets in bipolar disorder. Psychiatry Research 257, 327–337 (2017)

[106] Hu, V.W., Frank, B.C., Heine, S., Lee, N.H., Quackenbush, J.: Gene expression profiling of lymphoblastoid cell lines from monozygotic twins discordant in severity of autism reveals differential regulation of neurologically relevant genes. BMC genomics 7, 1–18 (2006)

[107] Scheinfeldt, L.B., Hodges, K., Pevsner, J., Berlin, D., Turan, N., Gerry, N.P.: Genetic and genomic stability across lymphoblastoid cell line expansions. BMC research notes 11, 1–5 (2018)

[108] Romanovsky, E., Choudhary, A., Akel, A.A., Stern, S.: Seeking convergence and divergence between autism and schizophrenia using genomic tools and patients’ neurons. bioRxiv, 2023–08 (2023)

[109] Stern, S., Zhang, L., Wang, M., Wright, R., Rosh, I., Hussein, Y., Stern, T., Choudhary, A., Tripathi, U., Reed, P., et al.: Monozygotic twins discordant for schizophrenia differ in maturation and synaptic transmission. Molecular Psychiatry, 1–15 (2024)

[110] Hussein, Y., Tripathi, U., Choudhary, A., Nayak, R., Peles, D., Rosh, I., Rabinski, T., Djamus, J., Vatine, G.D., Spiegel, R., et al.: Early maturation and hyperexcitability is a shared phenotype of cortical neurons derived from different asd-associated mutations. Translational Psychiatry 13(1), 246 (2023)

[111] Peykov, S., Berkel, S., Schoen, M., Weiss, K., Degenhardt, F., Strohmaier, J., Weiss, B., Proepper, C., Schratt, G., Nöthen, M., et al.: Identification and functional characterization of rare shank2 variants in schizophrenia. Molecular psychiatry 20(12), 1489–1498 (2015)

[112] Guilmatre, A., Huguet, G., Delorme, R., Bourgeron, T.: The emerging role of shank genes in neuropsychiatric disorders. Developmental neurobiology 74(2), 113–122 (2014)

[113] Chen, R.-S., Deng, T.-C., Garcia, T., Sellers, Z.M., Best, P.M.: Calcium channel *γ* subunits: a functionally diverse protein family. Cell biochemistry and biophysics 47, 178–186 (2007)

[114] Guan, F., Zhang, T., Liu, X., Han, W., Lin, H., Li, L., Chen, G., Li, T.: Evaluation of voltage-dependent calcium channel *γ* gene families identified several novel potential susceptible genes to schizophrenia. Scientific reports 6(1), 24914 (2016)

[115] Cunha, S.R., Mohler, P.J.: Ankyrin protein networks in membrane formation and stabilization. Journal of cellular and molecular medicine 13(11-12), 4364–4376 (2009)

[116] Wirgenes, K.V., Tesli, M., Inderhaug, E., Athanasiu, L., Agartz, I., Melle, I., Hughes, T., Andreassen, O.A., Djurovic, S.: Ank3 gene expression in bipolar disorder and schizophrenia. The British Journal of Psychiatry 205(3), 244–245 (2014)

[117] Falls, D.L.: Neuregulins: functions, forms, and signaling strategies. The EGF Receptor Family, 15–31 (2003)

[118] Meier, S., Strohmaier, J., Breuer, R., Mattheisen, M., Degenhardt, F., Mühleisen, T.W., Schulze, T.G., Nöthen, M.M., Cichon, S., Rietschel, M., et al.: Neuregulin 3 is associated with attention deficits in schizophrenia and bipolar disorder. International Journal of Neuropsychopharmacology 16(3), 549–556 (2013)

