## Supplementary material for "Predicting Suicide Risk in Bipolar Disorder patients from Lymphoblastoid Cell Lines genetic signatures": https://github.com/omveersharmanet/Predicting-Suicide-Risk-in-Bipolar-Disorder-patients.git

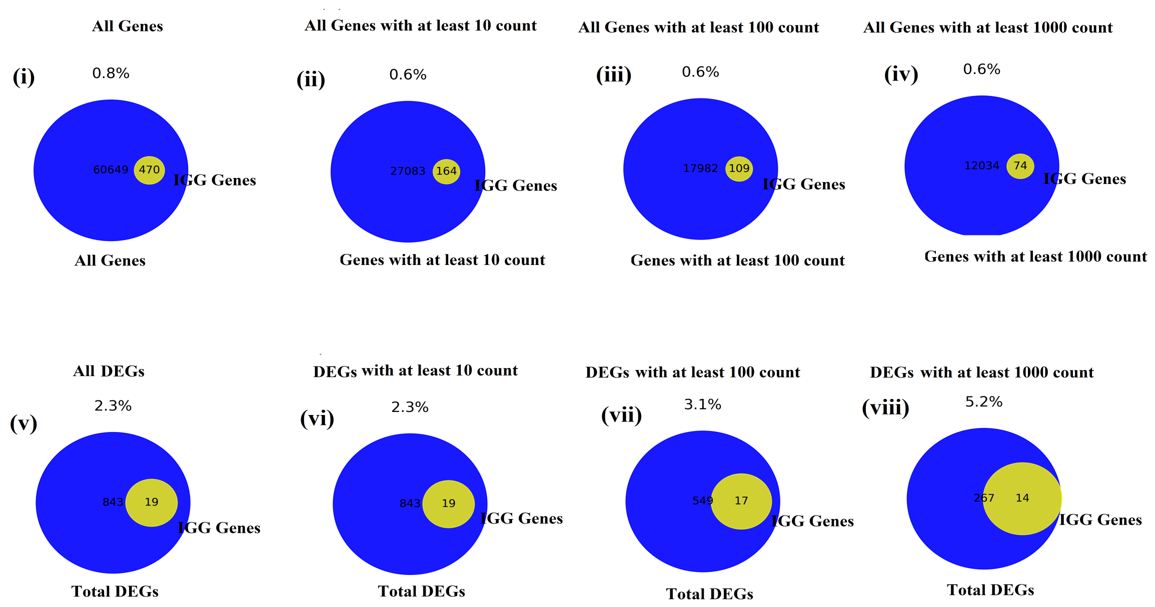


(a)

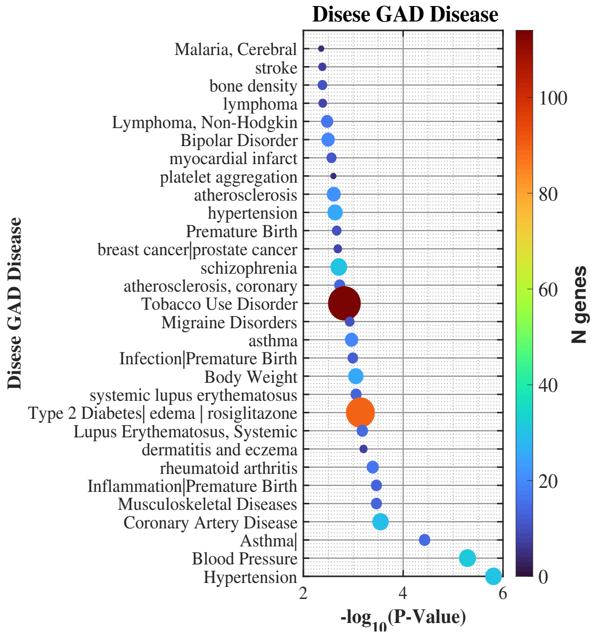


(b)

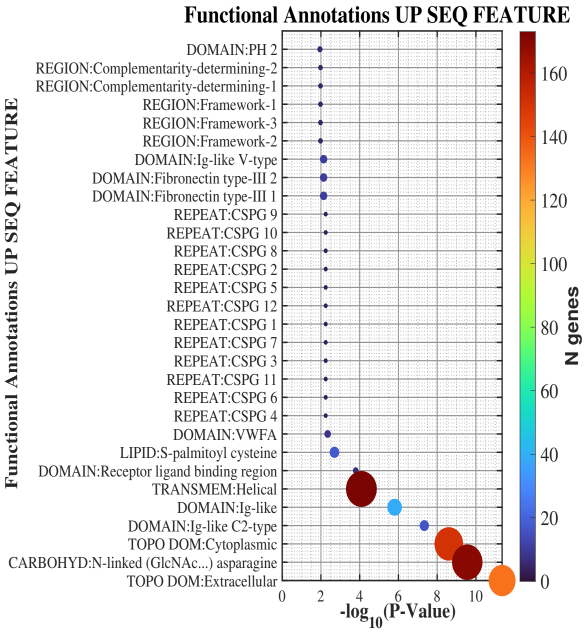


(c)

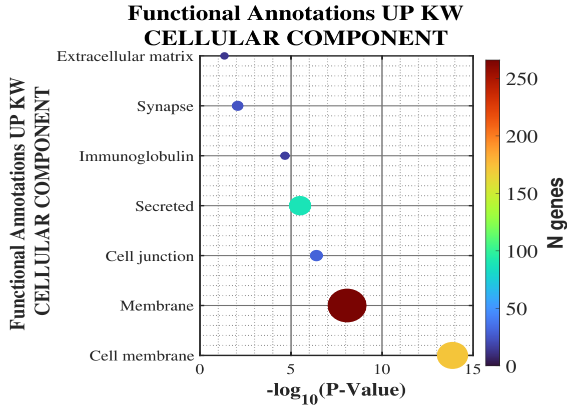


(d)

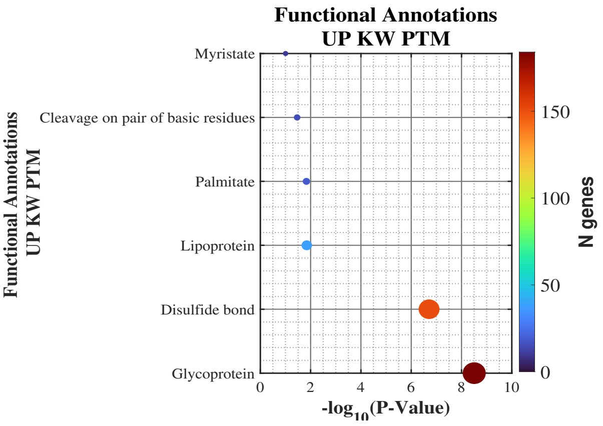


(e)

**Fig. 1**: Disease and functional annotation associations related to the DEGs using Disease DisGeNET and GAD Disease databases. (a) Percentage of IGG genes (yellow circle) are enriched in LCLs of BD patients who died by suicide, (i) Percentage of total IGG genes from the entire human gene list detected via RNA sequencing or genes with at least (ii) 10 or (iii)100 or (iv)1000 counts, (v) Percentage of total IGG genes in DEGs (between ‘SUICIDE’ and ‘NON-SUICIDE’) with at least (vi) 10 or (vii)100 or (viii)1000 counts. Enrichment of DEGs in (b) Disease GAD Disease. (c) Functional Annotations UP KW Cellular component. (d) Functional Annotations UP KW PTM. (e) Functional Annotations UP SEQ FEATURE.
